# CViT-ESP: Lightweight Pre-trained Vision Transformers for EEG-based Epileptic Seizure Prediction

**DOI:** 10.64898/2026.08.21.746341

**Authors:** Umair Mohammad, Paras Parani, Fahad Saeed

## Abstract

**Background and Objective:** Epileptic seizure prediction is a critical challenge requiring the discrimination of subtle preictal physiological changes from interictal brain activity. While deep learning has shown promise in this domain, existing models often face limitations due to small EEG datasets, high computational costs for training from scratch, and a lack of patient-independent generalizability. In this paper, we present a novel framework for EEG-based seizure prediction that leverages pre-trained Vision Transformers (ViTs) through custom architectural modifications and optimized re-training strategies.

**Methods:** Our primary contributions include:

- **CVIT-ESP:** A family of vision transformer architectures that replaces standard patch embedding layers with custom N-dimensional CNN stages to refine EEG representations.
- **ESPFormer:** A lightweight, custom-designed transformer specifically engineered to mitigate overfitting on limited-scale EEG datasets.

We identified optimal fine-tuning combinations for transformer blocks by devising a **heuristic search-space reduction strategy**, significantly reducing the training complexity. We validated our methods using the patient-independent **MLSPred-Bench**, involving 12 diverse benchmarks with varying seizure prediction horizons.

**Results:** Results demonstrate a clear progression in performance: while prior ResNet and vanilla Transformer models achieved an AUC-ROC of **69.0%**, our CVIT-ESP architectures achieved the highest performance with a maximum average AUC of **76.4%**.

**Conclusions:** These findings suggest that adapting pre-trained ViTs with domain-specific CNN front-ends and strategic fine-tuning offers a robust, generalizable, and resource-efficient path forward for clinical seizure prediction systems. Our code is available at: https://github.com/pcdslab/CVitEsp and https://github.com/pcdslab/ESPFormer

## Introduction

Epilepsy is the one of the costliest and second most common neurological disorder [1]. Epilepsy afflicts more than 3 million people in the US, with an estimated cost of US $24.5 Billion. The bulk of this health care spending is specific to seizure costs ($19 Billion) [2]. [3]. As per CDC statistics, adults in the US with active epilepsy, 41% reported poor health, 38% reported having a disability, and 42.4% (>1.1 million) reported having more than 1 seizure while taking anti-seizure medications [4]. These recurring, and unpredictable seizures significantly increase the risk of mild to severe injury, impact daily life by limiting social interactions/situations and physical activities – thereby reducing the quality of the life in general of children, adults and seniors. The growth in both inexpensive and noninvasive electroencephalography (EEG) sensors and accessible EEG data has been instrumental in accelerating the creation of computational approaches for seizure detection. While seizure detection is a valuable tool for safety monitoring, it addresses the consequences of a seizure after it has started. Seizure prediction, though still a major scientific and engineering challenge, holds the promise of changing the epilepsy experience by offering the chance to prevent or mitigate the seizure before it manifests, thereby leading to dramatic improvement in quality of life and patient autonomy. The ability to accurately predict seizures [5] could help patients regain some of the activities (driving [6], employment [7], leisurely activities[8]), reduced hospitalizations (Out of 9754 surveyed patients up to 58.8% of patients had ≥1 medically treated seizure[9]. [10]) and lower costs for families (costs per-patient-per-year ranged from $31,278 to $84,939). A system that could predict an incoming seizure for these patients and their families will improve individual’s perception of their physical, mental, and social well-being as it relates to their health. Notwithstanding the potential of a seizure prediction system to enhance the quality of life for millions, its success has been limited to date.

Predicting seizures is a considerably more complex problem than detecting them. It requires distinguishing the subtle pre-ictal phase from the interictal baseline, unlike detection which focuses on a more pronounced and readily identifiable difference between ictal and interictal activity [11]. Predicting a seizure may require holistic learning of the complex physiological changes that occur before, during, and after a seizure. The variability of physiological changes that accompany seizures are highly variable depending on types of seizures as well as across different individuals leading to subtle, indicative patterns in unequal distribution at various time points. Seizure detection and prediction machine learning (ML) models using EEG data have been proposed with mixed results when subject to independent evaluations. Reasons for inconsistent performance include both the problem complexity (variable seizures, non-smoothness, non-linearity, high-dimensionality, unimodality) and technical complexity (artifacts, limited annotated data, limited cohorts). In addition, inconsistent temporal windows, evaluation workflows prone to data-leakage, and onslaught of commercial products do not pass the filter of rigorous scientific testing. To facilitate development of seizure detection and prediction methods, multiple retroactive datasets with annotated seizures exist including the CHB-MIT and the Temple University Seizure Detection Corpus (TUSZ). To further facilitate researchers we recently introduced MLSPred-Bench [12] which makes ML-ready EEG data available suitable for developing predictive models. MLSPred-Bench, transforms raw EEG data into ML-ready datasets by automating the extraction of **preictal (pre-seizure)** and **interictal (normal)** windows across varying time horizons, which enables it to provide a consistent baseline for developing and evaluating deep learning models on a massive scale. Predicting a seizure remains an open research problem.

In this paper, we present the design and development of nine new architectures and strategies for EEG-based epileptic seizure prediction using pre-trained vision transformers (ViTs). Our primary contributions include the **CVIT-ESP** family, which redesigns the input stages of pre-trained ViTs with custom N-dimensional CNNs, and **VITESPOpt**, which utilizes a new heuristic search-space reduction strategy to find optimal fine-tuning combinations for transformer blocks. We also introduce data wrangling methods to adapt multidimensional EEG time-series data for image-based ViTs and develop **ESPFormer**, a custom lightweight transformer-based model designed specifically for the limited scale of EEG datasets. These models are validated using the patient-independent MLSPred-Bench, demonstrating improved generalizability, and a better balance between sensitivity and specificity compared to existing methods.

Our contributions include:

- **CVIT-ESP**: A new family of convolution-ViT architectures that replaces standard patch embedding layers with custom 1D or 2D CNN stages to refine EEG representations for transformer processing.
- **VITESPOpt**: A heuristic optimization strategy that reduces the exponential complexity of searching for effective fine-tuning combinations, resulting in substantial training time savings.
- **Data Wrangling**: Innovative methods to transform multidimensional EEG time-series into RGB-like image formats compatible with state-of-the-art pre-trained ViTs.
- **ESPFormer**: A lightweight, custom-designed transformer architecture specifically engineered to mitigate overfitting on limited EEG datasets.

We validate our proposed architectures—totaling nine new models and strategies—using the patient-independent MLSPred-Bench. The results from the MLSPred-Bench evaluation demonstrate that hybrid CNN-Vision Transformer (CViT-ESP) architectures consistently outperform (AUC scores of up to 76.4%) traditional 1D CNNs and ResNets, particularly on complex benchmarks with longer prediction horizons. A key finding is the superiority of SegFormer-based models over Swin-based ones, likely due to their use of learnable embeddings and GELU activation which better capture the chaotic nature of EEG signals. While the SegOpt model offers the highest sensitivity at 75.6%, making it ideal for patients with frequent seizures, the CSeg-ESP-1 model provides a more balanced profile with 10% higher specificity, effectively reducing the burden of false alarms. In addition our custom-designed lightweight transformer-based architecture called ESPFormer shows robust performance and is compared with results against those from the pre-trained ViTs with minimal fine-tuning, pre-trained ViTs with optimal fine-tuning and our custom-designed CViT-ESP. Key lessons from this result include that the increased data density in benchmarks with longer seizure prediction horizons (SPH) allows these transformer-based models to more effectively extract critical preictal biomarkers.

### Related work

In general, several deep learning (DL) architectures have been tested based on techniques including one or a combination of the following: convolutional neural network (CNN), long short-term memory (LSTM), transformers, etc. One of the prevailing architectures used in the literature is the combined CNN-LSTM including [13] which used it with neural architecture search (NAS). Other recent works have also used the CNN-LSTM approach including [14] and 3D CNN-LSTM [15]. The work of [16] used contrastive learning. Only recently, transformer architectures have been incorporated into CNN-LSTM models [17], [18] but mainly for feature extraction. The work of [17, p. 202] uses short-time Fourier Transform (SFTF) to extract temporal features followed by a transformer to extract spatio-temporal features whereas the work of [18] uses a ViT to extract spatial correlations. Classification is performed by a combination of CNN-LSTM modules. All of those methods have achieved a maximum sensitivity above 95% but are patient-specific, report results from the more restrictive CHB-MIT dataset and either do not apply leave-one-out cross-validation (LOOCV) or apply it incorrectly.

All of the above referenced works [13], [14], [15], [16, p. 20], [17, p. 202], [18] test the models using a very small-sized cohort, and perform \emph{only} patient-specific seizure prediction. Though a few works such as [19], [20] develop patient-independent models which are more generalizable, they also use a small cohort and apply CV in a way that causes data leakage. CHB-MIT is widely used due to its high quality but comprises data from only 23 pediatric subjects lowering the chances of generalizable models. Building large datasets to train all seizure types is challenging due to the obstacles in acquiring expert annotated data, prevailing variations in epilepsy and seizure types and the type of EEG systems, and challenges in acquiring data for comorbid factors. Further, applying LOOCV techniques to high-performing models such as [13] with a prediction sensitivity of 99.6% reduces their performance [21] to below 70% sensitivity. Moreover, the transformer-based architectures used do not take advantage of pre-trained architectures that provide a high-performance in other domains or propose an end-to-end predictive model.

Transformers were first introduced in [22] and have revolutionized the field of ML/AI in the generative AI era. The original transformer architecture comprises two blocks, an encoder and a decoder where only the encoder is used for classification tasks such as seizure prediction. Though originally proposed for natural language processing (NLP) tasks and used in large language models (LLMs) such as the Longformer [23] , transformers have been successfully adopted for other domains, notably for computer vision tasks with techniques such as the vision transformer (ViT) [24], [25], [26], [27]. Further, there exist several pre-trained ViT and LLM architectures with billions of parameters trained on petabytes of data which are available open-source. While recent literature and our results show that the use of transformers is the right direction for seizure prediction, EEG datasets are too small to train large architectures and the resource requirement is exhaustive for training from scratch.

Hence, there are three potential directions for seizure prediction. The first approach is to partially re-train some layers (referred to as fine-tuning) of the pre-trained models with domain adaptation of EEG data and this has shown to work from our previous results. The second approach is to keep some of the middle pre-trained layers while innovating at the initial layers and in the classifier head. This combines custom-modifications with fine-tuning and can be done either by adding initial layers suited to EEG data or modifying the initial layers of the pre-trained architectures to handle EEG data. The last approach is to develop custom-architectures as done in the literature but reducing the size and number of parameters such that the model is lightweight and less susceptible to overfitting on smaller EEG datasets. Approaches 1 and 2 are specifically of interest as these ViTs and LLMs are expected to be deployed on smartphones and can easily be leveraged for seizure prediction. In this work, we build upon our previous experiences by (1) introducing an effective way of searching best fine-tuning strategies for ViTs, (2) creating a new version of ViTs by adding innovative layers to handle EEG data and (3) providing a new custom-designed architecture based on the basic transformer architecture.

Recall that an epileptic seizure has 4 stages: the preictal stage which is the duration before a seizure, the ictal phase which includes the main symptoms, a post-ictal recovery stage followed by the interictal where the brain functions normally between 2 seizures. Seizure prediction is a challenging task which involves discrimination of preictal and interictal segments, as compared to only detection (ictal phases from the non-ictal phases). To use an existing pre-trained model for other domains, we solve two major problems, namely, (1) how to perform domain adaptation on EEG data for ViTs and optimizing the search strategy for best fine-tuning practices of pre-trained layers; (2) how to add innovative components to pre-trained architectures to make them more suitable for EEG data. Further, we validate the models using validation data from our benchmark MLSPred-Bench [28] which includes seizures from subjects that are disjoint from the training set. Hence, the models are more generalizable compared to those trained with patient-specific approaches. The outline of our work is listed below.

1. We propose a new family of **<u>c</u>**onvolution-**<u>ViT</u>**architectures for **<u>e</u>**pileptic **<u>s</u>**eizure **<u>p</u>**rediction (CViT-ESP). Particularly, CViT-ESP redesigns the first stage of the pretrained ViTs by implementing a custom-designed *N*-dimensional CNN to create CViT-ESP-*N*, where ViT is a placeholder for two distinct ViTs Swin or Seg (for SegFormer) and *N* indicates the type of CNN, either 1D or 2D. When designed CViT-ESP with both 1D and 2D CNNs, the approach results in four new architectures: CSwin-ESP-1 and CSeg-ESP-1, and CSwin-ESP-1 and CSwin-ESP-2.
2. We propose **<u>ViT</u>**s for **<u>e</u>**pileptic **<u>s</u>**eizure **<u>p</u>**rediction with **<u>opt</u>**imal fine-tuning strategies (ViTESPOpt). Once again, ViT is a placeholder for either Swin or Seg which results in two approaches SwinESPOpt and SegESPOpt (for brevity, we often refer to these as SwinOpt and SegOpt respectively throughout the manuscript). We propose a heuristic approach to optimize the search space for possible re-training strategies for multi-stage ViTs; specifically targeting ViTs that have an input stage, multiple transformer blocks followed by the classification stage.
3. We propose wrangling methods to transform multidimensional time-series EEG data into a format suitable for ViTs trained on images. Then, use the transformed data in conjunction with **<u>ViT</u>**s for **<u>e</u>**pileptic **<u>s</u>**eizure **<u>p</u>**rediction (ViTESP) which results in two models: SwinESP and SegESP. The approach closely follows our work in [29] where we quantified the seizure prediction results with only re-training the input and classification stages after the data wrangling pipeline. This represents the use of ViTs with minimal modifications and fine-tuning.
4. Lastly, we develop a new custom-designed lightweight transformer-based architecture called **<u>e</u>**pileptic **<u>s</u>**eizure **<u>p</u>**rediction trans**<u>former</u>** (ESPFormer) and compare the results against those from the pre-trained ViTs with minimal fine-tuning, pre-trained ViTs with optimal fine-tuning and our custom-designed CViT-ESP. Additionally, we also compare against our previously published architecture **<u>s</u>**eizure **<u>p</u>**rediction using **<u>E</u>**EG data with **<u>R</u>**esNets and **<u>T</u>**ransfer **<u>L</u>**earning (SPERTL).
5. We propose a strategy to evaluate continuous-time raw EEG data using several of our proposed DL architectures and combine it with a pipeline to make our results more interpretable. In total, we propose nine new architectures/strategies for seizure prediction with pre-trained ViTs in this article including: CSeg-ESP-1, CSwin-ESP-1, CSeg-ESP-2, CSwin-ESP-2, SwinOpt, SegOpt, SwinESP, SegESP and ESPFormer.

## Methods

### Datasets and Tools

Our recent tool MLSPred-Bench [28] leveraged the Temple University Seizure Detection Corpus (TUSZ) to derive patient-independent seizure prediction benchmarks. We used these benchmarks to validate and compare all architectures. Recall that seizure prediction requires discriminating between preictal and interictal segments which are not clinically annotated. Hence, MLSPred-Bench defined the seizure prediction horizon (SPH) to be the time from which the preictal samples are extracted and the seizure occur(rence) period (SOP) as a small gap duration before the start of a seizure that guarantees prediction ahead of time. Because various SHP and SOP values are proposed in the literature, multiple values for both *SPH* ∈ {2, 5, 15, 20} minutes and minutes *SOP* ∈ {1, 2, 5} were used to generate 12 benchmarks. The second dataset used is the CHB-MIT dataset which is widely used in the literature to validate seizure prediction models. The CHB-MIT dataset was mainly used to perform an evaluation of the generalization capabilities of our pre-trained ViTs on patient-specific data. Seven subjects were chosen that provided three or more seizures with gap times of more than four hours which reflects a research-based strategy [21].

The python programming language is used to develop, train and test all architectures including these specific libraries: NumPy for array manipulation and intermediate data storage, h5py and pandas to load ML-ready EEG datasets, the TensorFlow and PyTorch libraries in conjunction with the HuggingFace API to download pretrained models and fine-tune them on our dataset,Weights & Biases (wandb) to manage all training experiments and generate trained checkpoints, Scikit-learn to obtain performance metrics and the Matplotlib and Seaborn libraries to visualize the results. All other tasks are accomplished with built-in libraries available as part of the Python 3.12.3 or higher distribution.

### Overview of the proposed vision transformers (ViT)-based architectures

All methods developed in this work are based on the vision transformer (ViT), specifically, pre-trained ViTs which differ from traditional transformers due to their ability to process images. Typically, ViTs start with an input stage where large images are split into smaller fixed-size patches which are used to generate linear encodings and positional embeddings. This encoded data is then passed through a series of transformer blocks which may be followed by fully connected layers for the downstream task, especially for classification. In this work, we use two pre-trained ViTs: the Swin architecture [24], [25], [26] from which we have previously built SwinESP and SegFormer [27] from which we previously created SegESP. For SwinESP and SegESP, we overcame the challenge of making complex multi-channel EEG time-series data suitable for ViT-based processing with a sequence of augmentation steps. Overall in this paper, we discuss 10 different architectures that were developed and tested in five stages as described in Table 1. Out of the 10 models/strategies, six methods comprising stages four and five are newly designed and discussed for the first time in this paper.

**Table 1.** Architecture Descriptions. A description of how the 10 architectures compared in this paper were developed in 5 stages. The stage-wise development process will be justified by the results as shown in later sections.

| Arch. | Stage | Acronym | Name Description | Comments |
| --- | --- | --- | --- | --- |
| 1 | 1 | SPERTL | Seizure prediction with EEG and Transfer Learning | Referred to simply as ResNet sometimes in the manuscript |
| 2 | 2 | ESPFormer | Epileptic Seizure Prediction Transformer | A vanilla transformer customized to process EEG data for classification |
| 3 | 3 | SwinESP | Swin for Epileptic Seizure Prediction (ESP) | ViTESP: Fine-tuned ViTs for ESP. Swin and SegFormer were used with EEG data manipulation, a classifier head and fine-tuning with all transformer blocks frozen. |
| 4 |  | SegESP | SegFormer for ESP |  |
| 5 | 4 | SwinESPOpt (aka SwinOpt) | Swin for ESP with optimal search strategy | ViTESPOpt: ViT for ESP with optimal fine-tuning. SwinESP and SegESP are fine-tuned by only unfreezing a combination of transformer blocks as determined by our proposed heuristic strategy. |
| 6 |  | SegESPOpt (aka SegOpt) | SegESP with optimal search strategy |  |
| 7 | 5 | CSwin-ESP-1 | Convolution applied to Swin for ESP in 1D configuration | CViT-ESP-N: Convolution with ViT for ESP with $N$-d CNN. Instead of EEG data manipulation, we use a custom CNN architecture to extract embeddings for input to the first transformer block of SwinESP and SegESP. We only train the CNN block, the input stage of each CViT-ESP, and the classifier head without fine-tuning transformer blocks.. |
| 8 |  | CSeg-ESP-1 | Convolution applied to SegESP in 1D configuration |  |
| 9 |  | CSwin-ESP-2 | Convolution applied to Swin for ESP in 2D configuration |  |
| 10 |  | CSwin-ESP-2 | Convolution applied to SegESP in 2D configuration |  |

Four architectures belonging to the first 3 stages have been published previously with limited analysis/results with the first architecture based on the ResNet published as SPERTL [30], the second architecture as ESPFormer in [31] and the third and fourth architectures respectively referred to herein as SwinESP and SegESP and based on Swin and SegFormer published in [29]. The major contributions with respect to ESPFormer, SegESP and SwinESP in this paper include conducting comprehensive experiments across all MLSPred-Bench benchmarks for a comprehensive quantification of their performance. The two major contributions to model development in this paper include: (1) the development of an optimal search space reduction strategy for discovering the best retraining combinations of the transformer blocks which leads to SwinESPOpt (SwinOpt) and SegESPOpt (SegOpt) and (2) the development of a new architecture with an addition of custom-designed CNN modules leading to CVit-ESP. The ‘ViT’ in CViT-ESP is a placeholder for the base architecture (either Swin or Seg for SegFormer). Further, a digit ‘1’ or ‘2’ is appended to describe the implementation of the convolution operation in the CNN block which can be 1D (multiple 1D EEG channels) or 2D (2D convolution on multi-channel data). Hence, we end up with four distinct architectures: CSwin-ESP-1, CSeg-ESP-1, CSwin-ESP-2 and CSeg-ESP-2. We first describe CViT-ESP which fundamentally includes the convolutional block added to extract embeddings from multi-channel EEG data before combining with ViTs. This is followed by the description of the optimal re-training strategy with search space reduction that leads to SegESPOpt and SwinESPOpt For completeness, we summarily describe the SwinESP and SegESP architectures of [29, p. 202]. For self-containment, we also describe the ResNet and ESPFormer in the last subsection.

### CViT-ESP

Figure 1 illustrates the overall CViT-ESP architecture where we propose novel innovations to the existing Swin and SegFormer architectures to make the models compatible with and suited for classification of EEG data. Specifically, we add an additional custom-designed CNN stage that replaces the patch embedding layer which is part of the input stage of the original architecture. This CNN stage contains several convolutional layers, batch normalization, activation functions, and pooling layers which achieves two objectives: extracting additional spatial and temporal features before processing by the transformer stages and progressively compressing and refining the EEG data into a representation suitable for processing by the ViT stages. The CNN stage architecture is as follows: there are two convolutional - batch normalization (BN) layers, with CNN kernel being size 3 × 3 and BN based on standard scaling. This is followed by a max pooling layer that subsamples the output by two. The subsampled output then passes through another two convolution – BN layers followed by a second max pooling layer that further subsamples the output by two. This is followed by rectified linear unit (ReLU) activation and BN. After the last layer, the data is replicated to make it ready for processing with a ViT.

**Figure 1.**
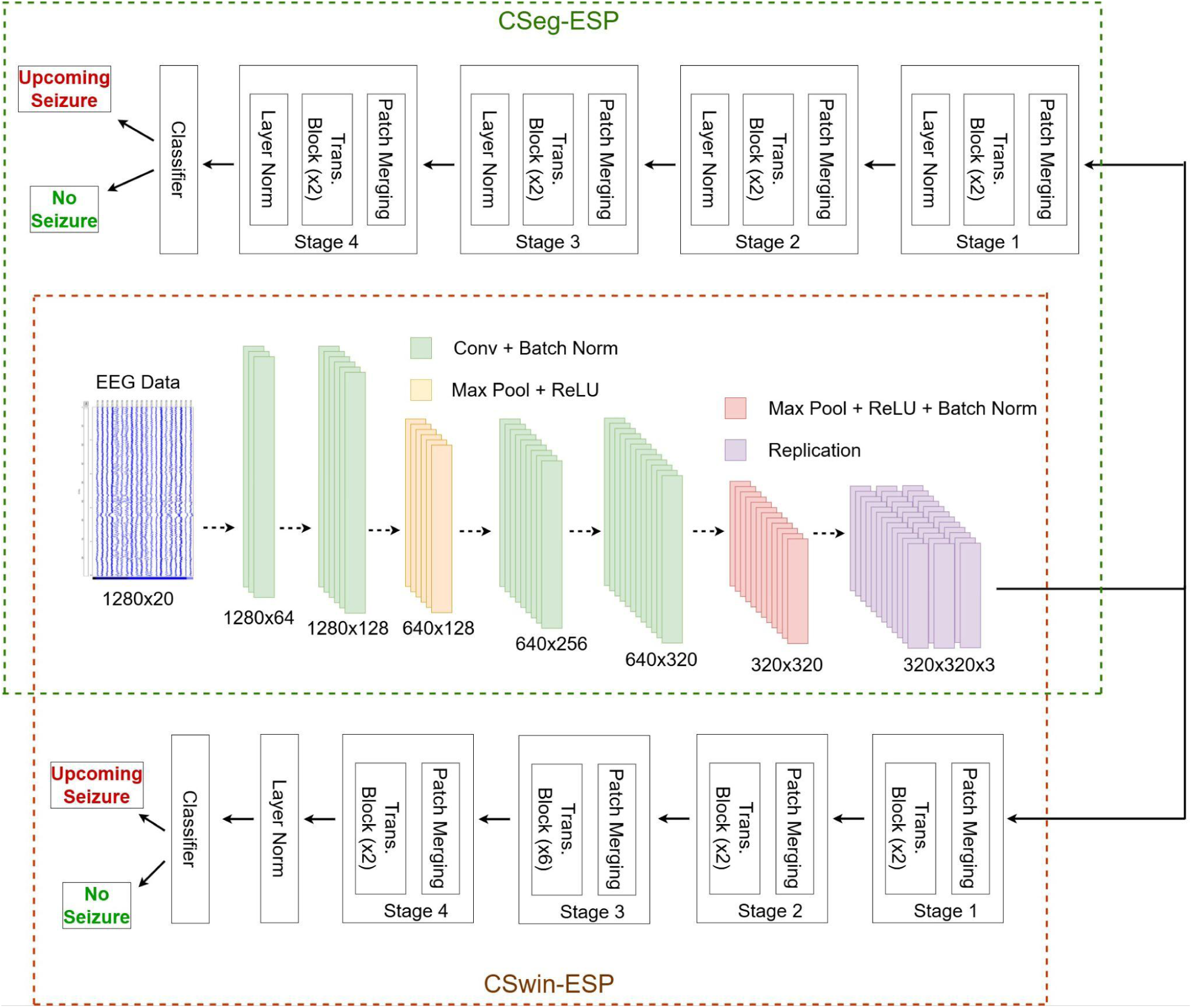
Depiction of the custom CNN added before the Swin/SegFormer Transformer blocks. The architecture is a simple two-stage CNN, each stage comprising two successive convolutional filters and batch normalization pairs, followed by max pooling and ReLU activation. The two convolutional blocks are followed by a batch normalization layer, the output of which is input to the embedding stage of the ViTs.

From the data processing perspective, the EEG data from MLSPred-Bench contains short-duration EEG segments from 20 channels of length 5 seconds sampled at 256 Hz and labeled either as preictal (label ‘1’) or interictal (label ‘0’). Hence, each input segment is of size 1280 × 20; this data is input to the CNN stage. In each stage of the convolution, we increase the representations progressively where the input from 20 channels is projected onto 64 channels by the first convolutional layer, the second layer outputs 128 representations, the third layer 256 and the last convolution layer outputs 320 representations. The first max-pooling layer after the second convolutional layer reduces the time-steps from 1280 to 640 to output a sequence of shape 640 × 128 and the second max pooling layer after the fourth convolution kernel further reduces it by half to generate a 320 × 320 square representation. This form can be replicated three times to feed the ViT that expects a color RGB-like image of size 320 × 320 × 3; this is a common method used to process grayscale images with CNNs designed for color images.

Recall that we propose two versions, one with 1D convolution kernels and the second with 2D convolution. The 1D format except 1280 × 1 time-series with the number of input channels set to 20. From the implementation perspective, each convolutional layer hence modifies the number of channels it outputs while maintaining the size of the sequence (i.e. the temporal steps). While the focus is on the extraction of local temporal features, the spatial integrity is maintained with this approach. The convolution kernel is set to a size of 3. In the second approach, we implemented a 2D convolutional variant where all operations mirror the 1D version but use their 2D counterparts – for example, a Conv2d layer with a kernel size of (3, 1) replaces the Conv1d layer. This modification was made to compare results and determine if the 2D approach offers any advantages, even though theoretically both operations are equivalent. Regardless, both approaches results in each segmented converted into a 320 × 320 representation, which we replicate to generate the 320 × 320 × 3 image-like representations for input to the patch merging layers of both ViTs. The CNN layer can be combined with SegESP to form CSeg-ESP as shown in the top-half of Figure 1, or with SwinESP to for CSwin-ESP as shown in the bottom-half.

While we discuss SegESP and SwinESP later for completeness, the overall process for CSeg-ESP adds four transformer stages to the CNN stage, each transformer stage comprising patch merging, two transformer blocks with attention – fully connected (FC) layers and layer normalization. Layer normalization differs slightly from BN in CNNs as it is applied at the level of features as opposed to being applied across the whole batch in BN. Anyways, the last transformer stage is followed by the classifier head. The architecture of Swin is similar with a few variations. The transformer stages do not include layer normalization and the third stage includes six transformer blocks in contrast to all other stages which have two transformers. The last layer is once again a classifier head consisting of FC layers and because this is a binary classification

### SwinESPOpt and SegESPOpt

While pre-trained ViTs have great potential, the exhaustive amount of resources required for fine-tuning necessitates a smart re-training strategy. Both pre-trained ViTs used in our experiments comprise six stages. In each ViT, the first two stages generate the patch embeddings followed by four stages of transformer blocks and a classifier head in the last stage. One approach is to directly apply the models to the data after transformation. However, this approach fails mainly because the classifier head is randomly initialized. Ablation studies in our previous work demonstrate that only re-training the classifier or the embedding stage also provides sub-optimal performance compared to re-training both stages 1 and 6, the embedding layer and the classifier head. Ideally, it would be best to fine-tune the complete model on all benchmarks, but this would require exhaustive resources and may not be feasible for smaller HPC systems. Further, fine-tuning the whole model on small datasets may lead to overfitting. In contrast, re-training or tuning a subset of transformer stages may be more prudent and also feasible. A few alternate approaches have been applied in areas such as personalization where only the last few personalization layers are re-trained after the cut layer. Other approaches also re-train the first few and the last few layers while keeping the layers in-between frozen. However, these techniques have only been tried for DL architectures based on convolutional layers. To deal with the size and exhaustive resource-requirements of ViTs, we propose a new heuristic strategy instead of selecting some layers to re-train based on an ad-hoc selection. Herein, we propose SwinESP optimized (SwinESPOpt) and SegESP optimized (SegESPOpt) which incorporate an optimal search strategy to find the best combination of transformer blocks for re-training.

### Search Space Reduction Strategy

Both pre-trained ViTs used in this work comprise six well-defined stages with four transformer stages, none of which are tuned in SwinESP and SegESP. In general, many ViT architectures contain well-defined transformer blocks. Let us formally define the brute-force search needed to discover the best combination of stages required for fine-tuning with the best performance. Consider an architecture that comprises *N* stages where only *k* are re-trained at a time. Then, the total number of possibilities for with *k* stages is given by:

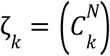

However, when we successively try re-training *k* = 0, 1, 2, …, *N* stages at a time, the total choices of for architectures with *N* > 2 can be expressed as followers:

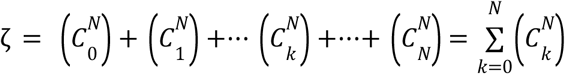

From the binomial theorem, we know that the number of combinations ζ can be given by *^N^* which represents an exponential complexity in terms of the number of experiments required. As an example, for SwinESP and SegESP where *N* = 6, this results in 64 combinations for a single architecture on a single dataset without any hyperparameter tuning. Given that we have 12 benchmarks and we test 2 pre-trained architectures, this represents 1536 choices of experiments. Further, the hyperparameters must be tuned for each approach to obtain the best predictive performance and hence, simply testing for a combination of 10 hyperparameters would result in an excess of 15,000 experiments. If we only consider *n* = 4 transformer stages out of *N* = 6 total stages, the number of experiments would still be greater than 3800. Since previous results indicate the necessity for re-training the embedding and classification phase, we focus our attention on the transformer stages and derive theoretical bounds on the complexity in terms of the number of experiments. Based on these derivations, we propose a theoretical heuristic algorithm and show how it can practically be applied to SwinESP and SegESP.

### Proposed Heuristic Algorithm (Theoretical Analysis)

Consider the total number of architectures to be trained as *L* architectures, for a total of *D* datasets, and tuned over a set of hyperparameters with cardinality *H*. In total, the cost of doing an exhaustive brute-force search to find the best fine-tuning strategy using all architectures, datasets and all hyperparameter sets in terms of the number of experiments would be:

*O_brute_* (*n*)=*LDH*2*^n^*

The objective of our approach is to reduce the number of possible combinations needed substantially. Due to the complexities of the data and underlying transformer-based architectures, it is difficult to analytically determine the most optimal re-training strategy. Hence, we propose a heuristic algorithm that uses the following three steps: combination reduction, hyperparameter tuning and best strategy selection. In the first step, instead of testing on all *D* datasets and *H* hyperparameter sets, we choose *D*’ out of *D* datasets as representative datasets and an initial set of hyperparameters to reduce the number of re-training combinations to:

*O*_1_ (*n*) = *LD*’2*^n^*

This approach is justified because either the datasets or the tasks for which the model is being trained may be similar. Further, current re-training strategies employed in the literature usually select the hyperparameters recommended by the developers of the model or start from the last set of hyperparameters on which the model converged for the original task.

In the second step, after enumerating the AUC scores on all architectures for the representative *D*’ datasets with 2*^n^* possible combinations, we choose *P* of the best ranked strategies to tune the hyperparameters. Here, the choice of best ranking can be purely objective, subjective or a combination of both and the choice of *P* is also flexible ranging from 1 to a maximum of 2*^n^*. An example of an objective methodology would be ranking all strategies based on the validation AUC score across all architectures, and then selecting the best *n* strategies. However, this can be combined with a subjective approach. Consider a model that comprises 4 transformer blocks. If the AUC score of re-training only stage 1 and 2 is marginally lower than re-training stages 1, 2, and 3, the second strategy can be skipped as it involves training an additional millions of parameters without considerable gain. With respect to the choice of *P*, choosing a small value such as 1 or even *n* may eliminate too many combinations that may work better for datasets not trained yet (recall that until this point we are using only *D*’ representative datasets). However, setting *P* too large such as on the order of *n*^3^ may raise the complexity too much without providing performance gains. Hence, we recommend *P* = *n* log *n*. In that case, to test all models with all *N* set of hyperparameters, the number of experiments needed for the second step are:

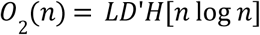

However, for complexity analysis, we will show the results for multiple cases of *P* including log(*n*), *n*, *n* log(*n*), *n*^2^ and *n*^3^ In that case, the complexity of the second stage would be *LD’HP* where *P* is a function of *n* given by *g*(*n*).

In the last step, we further reduce the number of re-training strategies by selecting the best hyperparameters for each architecture for each representative dataset and ranking them according to the validation AUC scores. The recommendation is to try at least close to *n* strategies at this stage (even if *P* was set to *n* in step 2, nearly all *n* strategies should then be used with the best hyperparameters discovered). All *L* architectures will now be trained all on *D* datasets using the best hyperparameters for each of the *n* final selected strategies. The number of experiments needed at this stage will be:

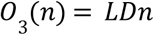

The heuristic algorithm’s complexity in terms of the total number of experiments can then be given by:

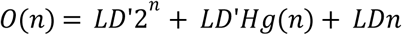

As noticeable, the choice of *P* = *g*(*n*) will have a significant impact on the overall number of experiments required. The drawback is that we still have an exponential term in step 1, but when testing a large number of architectures with many transformer blocks and on multiple datasets, steps 2 and 3 will offer significant reductions in complexity in terms of the number of experiments. Considering that each experiment represents training a DL model on a GPU and is of intractable time complexity, any method that can reduce the number of experiments by an order of magnitude would result in significant training time savings as demonstrated by our results.

### Implementation of SegOpt and SwinOpt

We apply the above proposed algorithm to tuning SwinESP and SegESP. In our work, we have *L* = 2 base architectures given by SwinESP and SegESP. When using MLSPred-Bench, *D* = 12 when each BM is considered a dataset. Hence, we set *D*’ = 2 and restrict the experiments for parameter tuning to two benchmarks, BM5 and BM8, and the justification is as follows. We notice the benchmarks are grouped by the SPH with BM1 – BM 3 using an SPH of two minutes, BM4 – B6 five minutes, BM7 – B9 fifteen minutes and BM10 – B12 thirty minutes. In each group, each successive benchmark uses an SOP of 1, 2 and 5 minutes respectively. Therefore, we opt for the middle benchmark to have a reasonable time in warning. Overall, the performance of BM11 is an outlier whereas BM1 – BM3 in general provide a low performance. Hence, we pick BM5 where the ViTs have shown to provide the best performance and BM8 where the ViTs provide a good performance but underperform other architectures. Further we optimized over a set of *H* = 10 hyperparameters using the weights and biases optimization tool. For step 2, we used *P* = 6 (exactly using *P* = *n* log *n* would have necessitated shortlisting 8 combinations) and for the final stage, we shortlisted 3 strategies (implying *n* − 1 combinations which is still of linear - *O*(*n*) - complexity) using a subjective approach. The justification for our choices is described in the results section.

### SwinESP and SegESP

Recall that SwinESP was adopted from Swin [24] which originally comprised six stages, an input stage that includes the patch partitioning and embedding layers followed by four transformer stages. We added a sixth classifier stage to create SwinESP. Further, we used a version of Swin [25] that modifies the attention mechanism by calculating the softmax output using a scaled cosine function version of the query and key values [26]. Initially, the model accepts data with an image-like representation of shape *H* × *W* × 3, and applies a patch partitioning kernel of size of 4 × 4 × 48. This results in patches of size *H*/4 × *W*/4 which are passed through an embedding layer that projects into space *C*, and *C* is a tunable parameter. Over the four transformer stages, the data is resampled as follows: *H*/4 × *W*/4 × *C* → *H*/8 × *W*/8 × 2*C* → *H*/16 × *W*/16 × 4*C* → *H*/32 × *W*/32 × 8*C*. In the classification stage, we perform 1D average pooling followed by FC layers and softmax activation.

SegESP was based on the Segformer [27] which was originally designed for semantic segmentation. As shown in the top half of Figure 1, this model also contains four transformer stages; each comprising both an encoder and a decoder. However, instead of positional embeddings, given an input sequence of feature maps *U_in_* from the self-attention modules, *FC*(.) representing the application FC layers and

*CNN* representing convolutions with 3 × 3 kernels, the model first obtains:

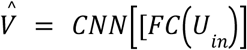

Then, the embedding representations are applying the Gaussian Error Linear Unit (*GELU*) activation function and FC layers as follows:

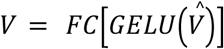

As mentioned earlier, each transformer stage contains patch merging with stage using a 7 × 7 kernel with stride length 4 and padding 3 × 3 whereas stages 3-5 use a 3 × 3 kernel with stride and padding 1 × 1.

The pretrained model originally used for SegESP came from the encoder-only network fine-tuned on ImageNet-1k given by the Nvidia/Mit-B0 checkpoint accepts images of shape 512 × 512 × 3. In contrast, SwinESP required data to be of shape 256 × 256 × 3. Recall that EEG data from MLSPred-Bench is represented by short 5-second duration segments of shape 1280 × 20 [28] which although similar in structure to an image is still not suitable for either ViT-based model. Therefore, we applied several augmentation steps based on simple replication that are applicable to SwinESP and SegESP, but also were used for SwinOpt and SegOpt in this work. For SwinESP, we further divided the segments into 1s duration and 2s for SegESP resulting in respective shapes of 256 × 20 and 512 × 20. Then, for SwinESP, along the channel axis we replicated the data 12 times to obtain a 256 × 240 representation, and added replications of randomly sampled 16 out of 20 channels which helped us obtain a 256 × 256 representation. A similar strategy was applied for SegESP where we replicated each channel 25 times to obtain a 512 × 500 representation, and added 12 channels selected randomly to generate a 512 × 512 representation. For both architectures, we replicated each 2D representation 3 times to obtain an RGB-like image of shapes 256 × 256 × 3 and 512 × 512 × 3 respectively for SwinESP and SegESP. This approach was also applied for SwinOpt and SegOpt. For CViT-ESP, we used checkpoints that accept 320 × 320 × 3 image-like representations.

We also add a classifier head that is randomly initialized to perform binary classification for SwinESP and SegESP which were fine-tuned using the following following hyperparameters: batch size of 128, 5 epochs, LR set to 1*e* – 3, a warmup ratio of 0.1 and 1% weight decay. The warm-up ratio is the ratio of training steps to the total steps needed to reach the pre-set learning rate, which is 1*e* – 3 in this case. For SwinOpt and SegOpt, we started with these hyperparameters but did additional tuning to generate the best hyperparameter set for various strategies, benchmarks and both architectures.

### ESPFormer (based on the transformer) and SPERTL (based on the ResNet)

For completeness, we cover the design of our prior architectures respectively based on the ResNet (SPERTL) and the vanilla transformer (ESPFormer). ESPFormer is novel in the aspect of visualizing each EEG sequence from 20 channels as a “sentence” with 1280 “words”. We modify the embedding stage so that the data is transformed from a 20-dimensional space to a 64-dimensional space, while maintaining the sequence length. Given an original sequence *X* ∊ *ℝ*^20×1280^, the new sequence will be:

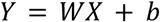

*W* ∈ ℝ^64×20^ is a trainable weight matrix and *b* is the bias vector. We add an additional *cls* token for classification by appending *cls^T^* ([.]*^T^* is the transpose operation) to the start of the sequence *Y* to create an aggregated representation *̂Y* ∊ *ℝ*^64×1281^for classification as follows:

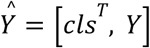

To glean the positional information, we use learnable positional embeddings *V* ∈ ℝ^64×1281^ to get

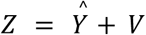

This sequence *Z* ∈ ℝ^64×1281^ contains both positional information and incorporates the classification token and is passed through two successive transformer blocks. Each transformer block consists of a single multi-headed attention module with four attention heads (to extract the temporal relationships) followed by normalization, an FC network and another normalization layer. To stabilize learning, each transformer block also contains a residual skip connection and uses GELU activation. After the second transformer block, we have the classification stage that contains an FC layer followed by ReLU activation and a second FC layer with sigmoid activation to generate the output probability. The SPERTL architecture is based on the residual neural network (ResNet) and features an input convolutional block followed by 4 residual blocks. Each convolutional block contains 3 × 1 convolution, BN, dropout (set to 0.75 for MLSPred-Bench) and 2 × 1 max pooling with ReLU activation. Each residual block contains 2 convolutional blocks; the skip connection in each residual block is located right after the convolution kernel of its second convolutional block. This differentiates SPERTL from a regular ResNet which typically has the skip connection between two convolutional blocks. The last residual block is followed by two FC layers and a softmax layer for classification.

## Results

### Results Overview

Overall, a comparison of the validation AUC scores indicates that there is a clear progression of improvement. Table 2 displays the AUC scores achieved for all architectures discussed in this paper including the six newly proposed methods CSwin-ESP-1, CSeg-ESP-1, CSwin-ESP-2, CSeg-ESP-2, SwinOpt and SegOpt and the previously published methods SwinESP, SegESP, ESPFormer and SPERTL In general, the CViT-ESP architectures provided the best average AUC scores across all benchmarks (BMs) with a minimum of 72.7% with CSwinESP-2 and a maximum of 76.4% with CSeg-ESP-1. In general, all CVit-ESP methods outperformed the other methods in terms of average AUC scores across all (BMs) with the exception of SegOpt with its AUC score of 74.5% beating CSwin-ESP-2 with an AUC score of 74.2%. Further, it was observed that there was a clear progression of AUC scores as our model development evolved from using simple transfer learning with ResNet to the custom-designed CNN-ViT architecture. For example, SPERTL and ESPFormer both provided an average AUC of 69.0%; ViTESP improved that to 69.7% with SegESP, though SwinESP had a worse AUC of 68.5%. By employing optimal fine-tuning strategies with ViTESPOpt, the AUC improved to 72.2% with SegOpt. SwinOpt also had a higher average AUC of 70.8% across all BMs though it was lower than SegOpt. However, at 72.2%, the average AUC of SegOpt was still bettered by all CViT-ESP approaches.

**Table 2.** AUC scores. A comparison AUC scores across all benchmarks for each model .

| BM | SPERT<br>L | ESPFo<br>rmer | SwinE<br>SP | SegES<br>P | SwinO<br>pt | SegOpt | CSwin-ESP-1 | CSeg--<br>ESP-1 | CSwin-ESP-2 | CSeg-ESP-2 | Avg. |
| --- | --- | --- | --- | --- | --- | --- | --- | --- | --- | --- | --- |
| BM1 | 66.8% | **79.8%** | 69.9% | 76.3% | 73.3% | 76.9% | 73.8% | 77.0% | 75.5% | 78.6% | 75.8% |
| BM2 | 66.5% | 77.6% | 72.6% | 76.2% | 77.8% | 76.7% | 75.3% | 77.3% | 75.1% | **80.3%** | 76.5% |
| BM3 | 69.4% | 59.9% | 73.3% | 76.1% | **76.2%** | 75.0% | 70.8% | 72.1% | 72.8% | 73.9% | 72.2% |
| BM4 | 63.2% | **82.7%** | 73.1% | 76.7% | 79.2% | 78.1% | 76.2% | 75.9% | 76.9% | 78.2% | 77.6% |
| BM5 | 62.4% | **80.4%** | 71.8% | 76.1% | 73.4% | 76.2% | 72.5% | 71.0% | 73.6% | 74.0% | 72.1% |
| BM6 | 64.0% | 74.4% | 73.7% | 77.0% | **79.1%** | 78.3% | 72.4% | 77.2% | 69.7% | 69.2% | 74.6% |
| BM7 | 81.1% | 61.6% | 73.0% | 71.9% | 73.9% | 80.8% | 78.9% | **81.1%** | 75.4% | 78.6% | 75.1% |
| BM8 | 83.6% | 61.8% | 71.4% | 72.0% | 76.1% | **83.8%** | 80.6% | 82.9% | 74.5% | 78.7% | 72.4% |
| BM9 | 79.2% | 59.7% | 74.2% | 71.2% | 75.5% | 77.3% | 81.3% | 81.7% | 77.5% | **81.9%** | 75.7% |
| BM10 | 56.4% | 53.4% | 43.6% | 41.0% | 48.4% | 49.4% | 72.8% | 68.0% | 70.5% | **73.1%** | 61.4% |
| BM11 | 80.5% | 83.0% | 80.2% | 78.7% | 84.0% | 88.2% | 91.8% | **93.7%** | 88.7% | 89.0% | 82.3% |
| BM12 | 50.8% | 53.5% | 45.0% | 42.8% | 55.2% | 53.2% | 69.8% | **71.4%** | 60.3% | 58.1% | 56.6% |
| Avg. | 69.0% | 69.0% | 68.5% | 69.7% | 72.7% | 74.5% | 76.4% | **77.4%** | 74.2% | 76.1% | 72.7% |

Tables 3 and 4 provide the other evaluation metrics including the accuracy, sensitivity and specificity for all ten architectures. Table 3 provides the metrics for SPERTL, SwinESP and SegESP, and SwinOpt and SegOpt whereas Table 4 illustrates the performance of ESPFormer, and all four CViT-ESP architectures. What distinguishes the architectures of Table 3 from Table 4 is that for the architectures of Table 3, we mainly focused on re-training/fine-tuning strategies with minimal modifications whereas Table 4 represents architectures where we added custom-designed layers. It is observable that SegOpt provides the highest metrics among SPERTL, ESPFormer, ViTESP and ViTESPOpt models with a validation accuracy of 69.8%, a prediction sensitivity of 75.6% and a specificity of 63.4%. SwinOpt provides a comparable sensitivity of 75.6% but has a lower specificity of 60.0%, which brings down its accuracy to 68.0%. Whereas ESPFormer, SwinESP and SegESP all provide respectable accuracies of 65.9%, 64.3% and 65.0% respectively, SPERTL had the lowest accuracy of 53.8%. This was mainly because of its low specificity of 39.9% as the sensitivity was a respectable 67.8%.

**Table 3.** Sensitivity, Accuracy and Specificity. Metrics for SPERTL, Swin/SegESP, Swin/SegOpt.

| Metri-<br>cs &<br>BMs | SPERTL |  |  | SwinESP |  |  | SegESP |  |  | SwinOpt |  |  | SegOpt |  |  |
| --- | --- | --- | --- | --- | --- | --- | --- | --- | --- | --- | --- | --- | --- | --- | --- |
|  | Acc | Sen | Spe | Acc | Sen | Spe | Acc | Sen | Spe | Acc | Sen | Spe | Acc | Sen | Spe |
| 1 | 61.8% | 90.8% | 32.9% | 66.4% | 75.5% | 57.3% | 69.7% | 66.5% | 72.8% | 69.2% | 88.5% | 50.0% | 74.3% | 87.4% | 61.3% |
| 2 | 50.0% | 99.7% | 0.4% | 68.2% | 66.8% | 69.6% | 70.7% | 73.9% | 67.5% | 71.8% | 87.4% | 56.2% | 73.3% | 80.2% | 66.4% |
| 3 | 55.1% | 98.4% | 11.7% | 68.5% | 76.3% | 60.7% | 68.6% | 69.6% | 67.8% | 68.9% | 81.2% | 56.6% | 70.3% | 77.8% | 62.7% |
| 4 | 57.8% | 63.2% | 52.4% | 65.7% | 71.7% | 59.6% | 67.5% | 73.5% | 61.5% | 73.2% | 79.4% | 67.1% | 75.6% | 78.1% | 73.0% |
| 5 | 57.4% | 29.1% | 85.7% | 67.2% | 73.0% | 61.4% | 68.3% | 73.8% | 62.7% | 69.2% | 71.0% | 67.5% | 70.1% | 76.4% | 63.8% |
| 6 | 60.0% | 22.9% | 97.1% | 67.7% | 75.0% | 60.5% | 70.3% | 76.1% | 64.5% | 72.4% | 81.4% | 63.5% | 72.1% | 72.6% | 71.6% |
| 7 | 53.0% | 7.0% | 99.0% | 67.3% | 76.2% | 58.4% | 67.0% | 75.2% | 58.8% | 68.4% | 77.4% | 59.4% | 74.3% | 74.0% | 74.6% |
| 8 | 51.0% | 2.6% | 99.3% | 65.8% | 76.2% | 55.4% | 65.3% | 71.0% | 59.4% | 69.2% | 81.6% | 56.9% | 68.7% | 89.0% | 48.5% |
| 9 | 49.9% | 99.6% | 0.2% | 67.2% | 75.3% | 59.1% | 64.7% | 69.4% | 59.9% | 69.5% | 80.7% | 58.2% | 70.3% | 77.9% | 62.7% |
| 10 | 50.0% | 100% | 0.0% | 48.4% | 61.6% | 35.3% | 46.4% | 66.8% | 26.0% | 53.0% | 66.7% | 39.3% | 53.6% | 68.0% | 39.1% |
| 11 | 50.0% | 100% | 0.0% | 70.0% | 63.9% | 76.1% | 71.2% | 64.9% | 77.5% | 76.8% | 59.6% | 82.5% | 80.1% | 69.6% | 83.5% |
| 12 | 50.0% | 100% | 0.0% | 48.9% | 61.1% | 36.7% | 50.1% | 63.2% | 37.1% | 54.1% | 45.1% | 63.2% | 54.7% | 55.8% | 53.6% |
| Avg. | 53.8% | 67.8% | 39.9% | 64.3% | 71.0% | 57.5% | 65.0% | 70.3% | 59.6% | 68.0% | 75.0% | 60.0% | **69.8%** | **75.6%** | **63.4%** |

**Table 4.** Sensitivity, Accuracy and Specificity. Performance of ESPFormer and CSwin/Seg-ESP-1/2.

| Metrics & BMs | ESPFormer |  |  | CSwin-ESP-1 |  |  | Cseg-ESP-1 |  |  | CSwin-ESP-2 |  |  | Cseg-ESP-2 |  |  |
| --- | --- | --- | --- | --- | --- | --- | --- | --- | --- | --- | --- | --- | --- | --- | --- |
|  | Acc | Sen | Spe | Acc | Sen | Spe | Acc | Sen | Spe | Acc | Sen | Spe | Acc | Sen | Spe |
| 1 | 74.3% | 80.4% | 68.2% | 66.6% | 70.2% | 62.9% | 72.6% | 85.8% | 59.3% | 68.0% | 76.3% | 59.8% | 73.8% | 85.4% | 62.1% |
| 2 | 72.8% | 78.7% | 67.0% | 70.5% | 83.7% | 57.2% | 71.9% | 87.1% | 56.8% | 69.4% | 79.9% | 59.0% | 73.6% | 88.7% | 58.6% |
| 3 | 62.4% | 75.6% | 49.1% | 65.5% | 86.1% | 45.0% | 66.2% | 66.5% | 65.8% | 65.9% | 70.8% | 61.1% | 66.3% | 77.9% | 54.6% |
| 4 | 75.5% | 81.5% | 69.5% | 73.3% | 76.6% | 70.1% | 70.5% | 78.0% | 63.1% | 71.3% | 76.4% | 66.3% | 73.1% | 80.8% | 65.4% |
| 5 | 74.0% | 80.5% | 67.5% | 66.7% | 75.9% | 57.5% | 69.3% | 74.7% | 63.9% | 68.7% | 72.0% | 65.4% | 68.7% | 74.6% | 62.9% |
| 6 | 68.6% | 75.5% | 61.6% | 65.6% | 65.1% | 66.0% | 72.0% | 67.5% | 76.4% | 66.4% | 65.0% | 67.8% | 65.8% | 59.1% | 72.5% |
| 7 | 60.5% | 68.2% | 52.7% | 73.7% | 63.8% | 83.6% | 74.7% | 66.0% | 83.4% | 70.1% | 72.3% | 67.9% | 71.1% | 68.3% | 73.9% |
| 8 | 60.8% | 69.8% | 51.7% | 74.5% | 65.5% | 83.4% | 74.9% | 68.5% | 81.3% | 71.4% | 61.3% | 81.4% | 70.8% | 50.8% | 90.8% |
| 9 | 57.8% | 65.4% | 50.3% | 72.4% | 64.3% | 80.5% | 72.3% | 64.5% | 80.1% | 67.9% | 61.8% | 73.9% | 71.6% | 73.5% | 69.8% |
| 10 | 53.3% | 30.7% | 75.8% | 70.9% | 64.3% | 77.5% | 68.7% | 56.5% | 81.0% | 57.7% | 26.3% | 89.0% | 67.8% | 57.0% | 78.6% |
| 11 | 76.6% | 20.1% | 95.4% | 85.1% | 70.2% | 90.1% | 87.5% | 70.9% | 93.1% | 80.5% | 58.4% | 87.9% | 80.6% | 49.3% | 91.0% |
| 12 | 54.0% | 23.0% | 84.9% | 68.0% | 61.0% | 75.1% | 69.9% | 63.4% | 76.4% | 58.7% | 47.9% | 69.4% | 56.8% | 41.2% | 72.5% |
| Avg. | 65.9% | 62.5% | 66.1% | 71.1% | 70.5% | 70.8% | **72.5%** | **70.8%** | **73.4%** | 68.0% | 64.0% | 70.8% | 70.0% | 67.2% | 71.1% |

SwinESP and SegESP without fine-tuning the transformer blocks have good respective sensitivities of 71.0% and 70.3% but have lower specificities of 57.5% and 59.6% respectively. Because ESPFormer has a relatively higher average specificity of 66.1% across all BMs, it provides a higher accuracy despite having a lower average AUC score. However, when the CViT-ESP architectures are considered, they outperform the other methods in most metrics with CSeg-ESP-1 providing the highest accuracy of 72.%, the highest sensitivity of 70.8% and the highest specificity of 73.4% as shown in Table 4. When fine-tuned models are considered, SegOpt beats CSeg-ESP-1 in the sensitivity metric with a value of 75.6% compared to 73.4%. However, CSeg-ESP-1 beats all other models in terms of AUC, accuracy and specificity. Before we delve into a detailed analysis of CViT-ESP results, we identify the best models from stages 3 – 5 of our model development process. Recall that SPERTL is the only model from stage 1 and similarly ESPFormer from stage 2. However, SegESP is the best performing compared to SwinESP from stage 3, SegOpt is better compared to SwinOpt and CSeg-ESP-1 provides the highest performance metrics among CSwin-ESP-1, CSeg-ESP-1, CSwin-ESP-2 and CSeg-ESP-2. Hence, we compare the best model of each stage.

Figure 2 shows the average performance of the best architecture from each stage across all benchmarks in terms of the prediction accuracy, sensitivity and specificity. In general, it can be observed that CSeg-ESP-1 provides the highest accuracy and specificity of approximately 72% and 74%, while it has a sensitivity of about 71%. The highest average sensitivity of around 76% across all benchmarks is provided by SegESPOpt, but it has a much lower specificity of about 64% leading to an accuracy around 69%. A good balance between sensitivity and specificity is important as a perfect sensitivity score may simply imply that all EEG segments are labeled as preictal. This occurs with our first model based on the ResNet across multiple benchmarks (BMs) including benchmark 1 (BM2), and BM9 - BM12. Therefore, it consistently provides the lowest accuracy. In contrast, the ESPFormer offered a better balance between both. Further, it was able to provide a comparable average accuracy to SegESP of around 65%. After executing the proposed search space strategy, the optimal retraining combinations of SegESPOpt are able to improve the accuracy to approximately 69%, which is further improved by CSeg-ESP-1 to 72%.

**Figure 2.**
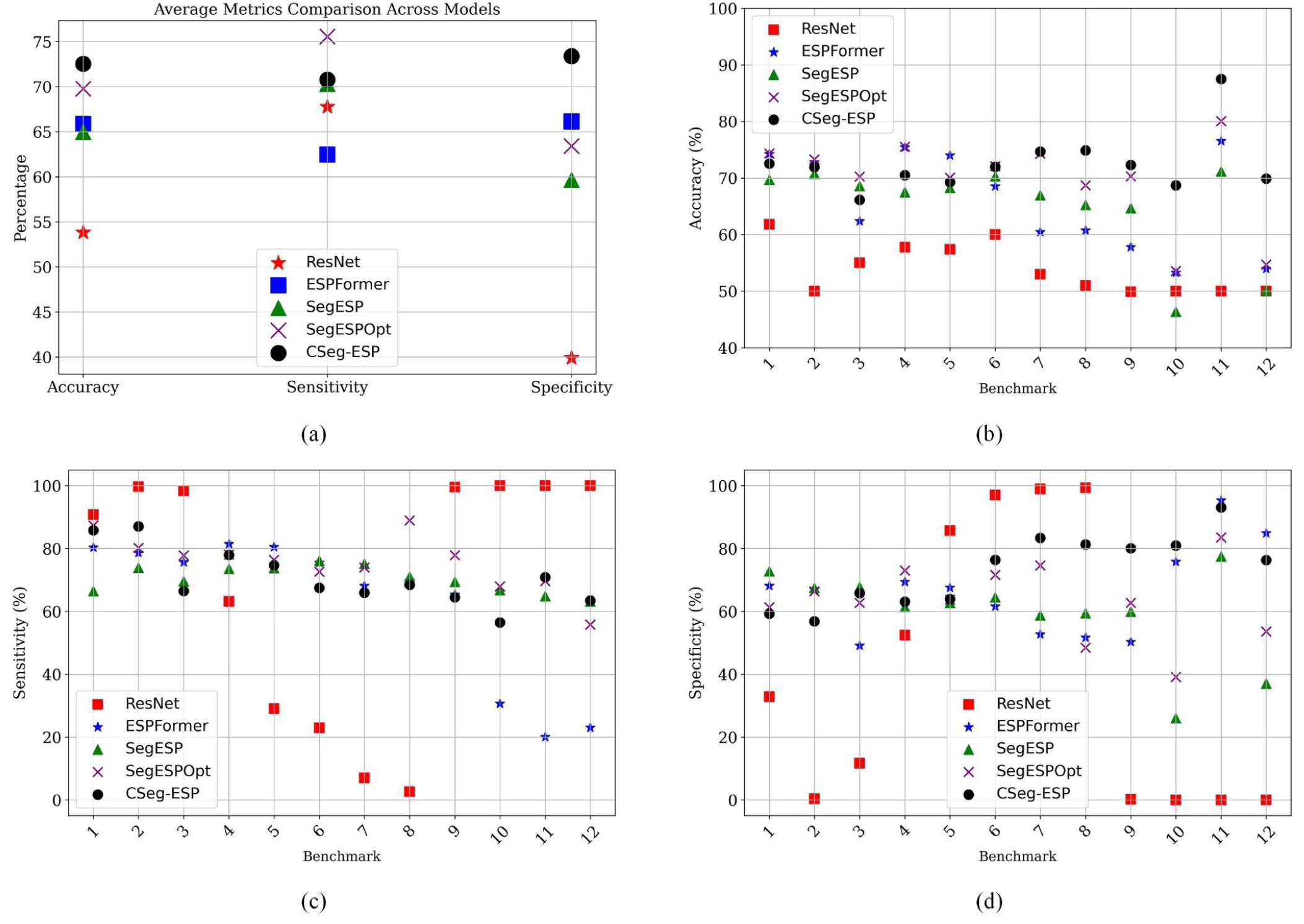
Average accuracy, sensitivity and specificity results. (a) Average of all three metrics for the best of each architecture type. (b) Average accuracy for each architecture.

### Results from CViT-ESP

Although CSeg-ESP-2 provided a competitive AUC score of 76.1%, the 1D versions of CViT-ESP outperformed the 2D versions where CSeg-ESP-1 had a higher average AUC of 77.4%, and CSwin-ESP-1 with an AUC of 76.4% and beat CSwin-ESP-2 which had an AUC of 74.2% as observable from Table 2. Figure 5 illustrates the ROC curves for all CViT-ESP architectures developed in this work. In general, after observing the curves, it is clear that adding a custom CNN block significantly bumps the ROC curves which increases the AUC scores. Particular improvements are noticed in BM10 - BM12. As Figure 6 shows, other architectures fail for these BMs, specifically, BM10 and BM12. Noticeable is the performance of CSeg-ESP-1 and CSwin-ESP-1 which have a significantly better ROC profile for BM12 and an improved AUC performance for BM12. This is reinforced by the average AUC scores in Table 2 where CSeg-ESP-1 provides an average AUC score of about 77.4% across all BMs and CSwin-ESP-1 provides an AUC score of 76.4%. This represents improvements of 5.6% and 4.2% compared to SwinOpt and SegOpt. Compared to the best performing model from stage four (SegOpt), these represent respective improvements of 1.4% and 4.2%.

**Figure 3.**
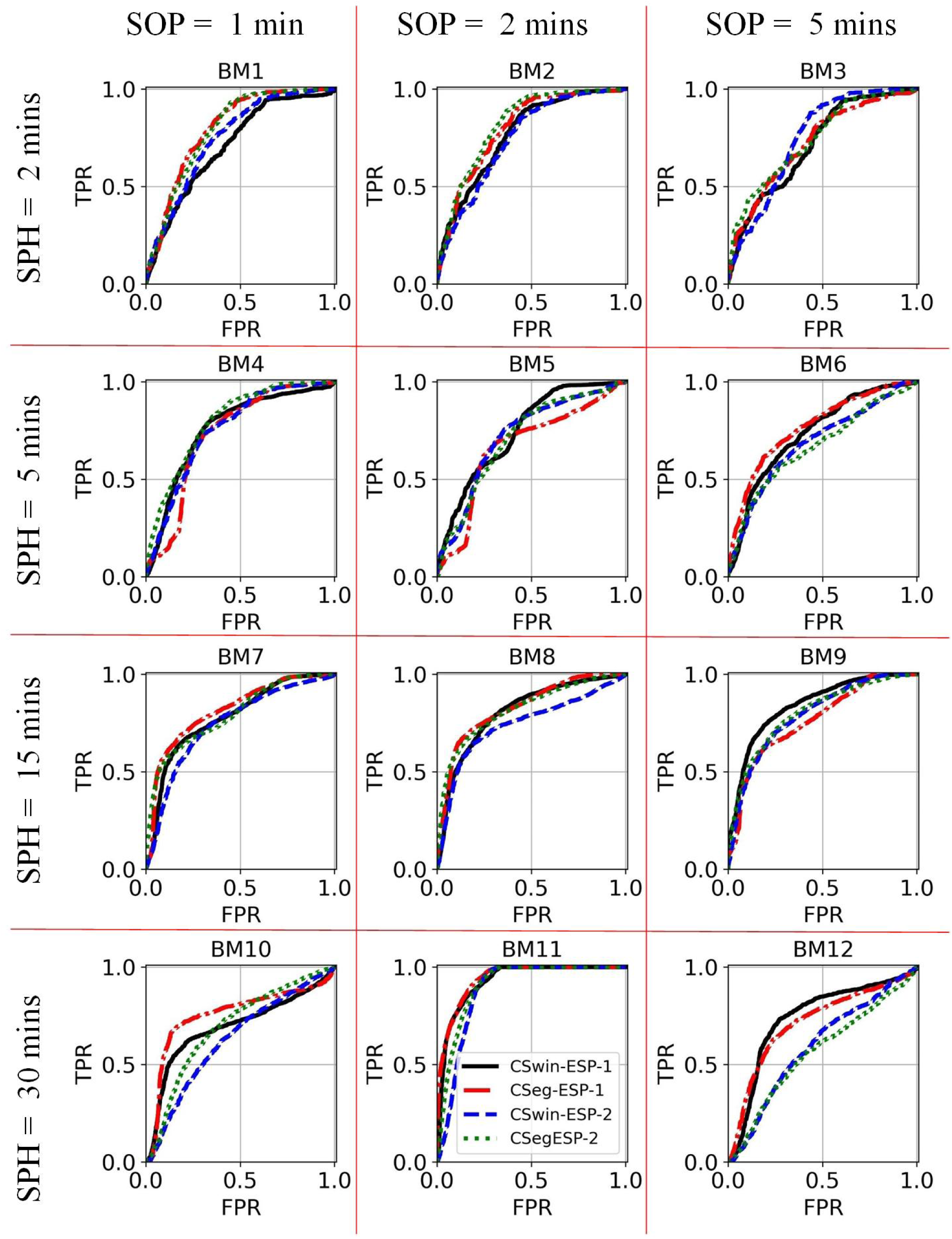
ROC Curves for custom modified architectures. The ROC curves for each benchmark for CSiwn-ESP-1, CSeg-ESP-1, CSiwn-ESP-2 and CSeg-ESP--2.

**Figure 4.**
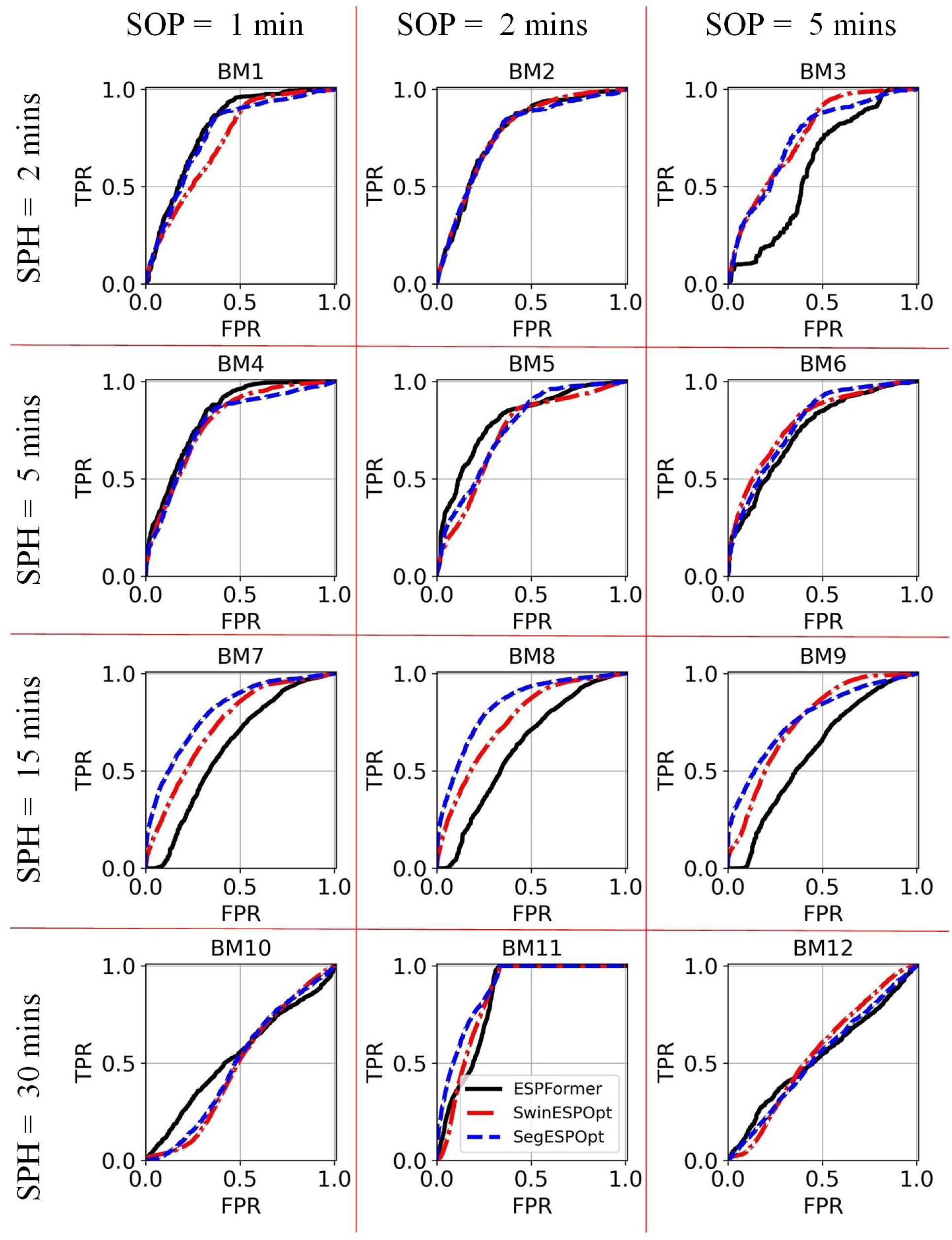
ROC Curves for ESPFormer compared to optimized Swin and SegFormer. The ROC curves for each benchmark for ESPSwinOpt, ESPSegOpt and ESPFormer.

**Figure 5.**
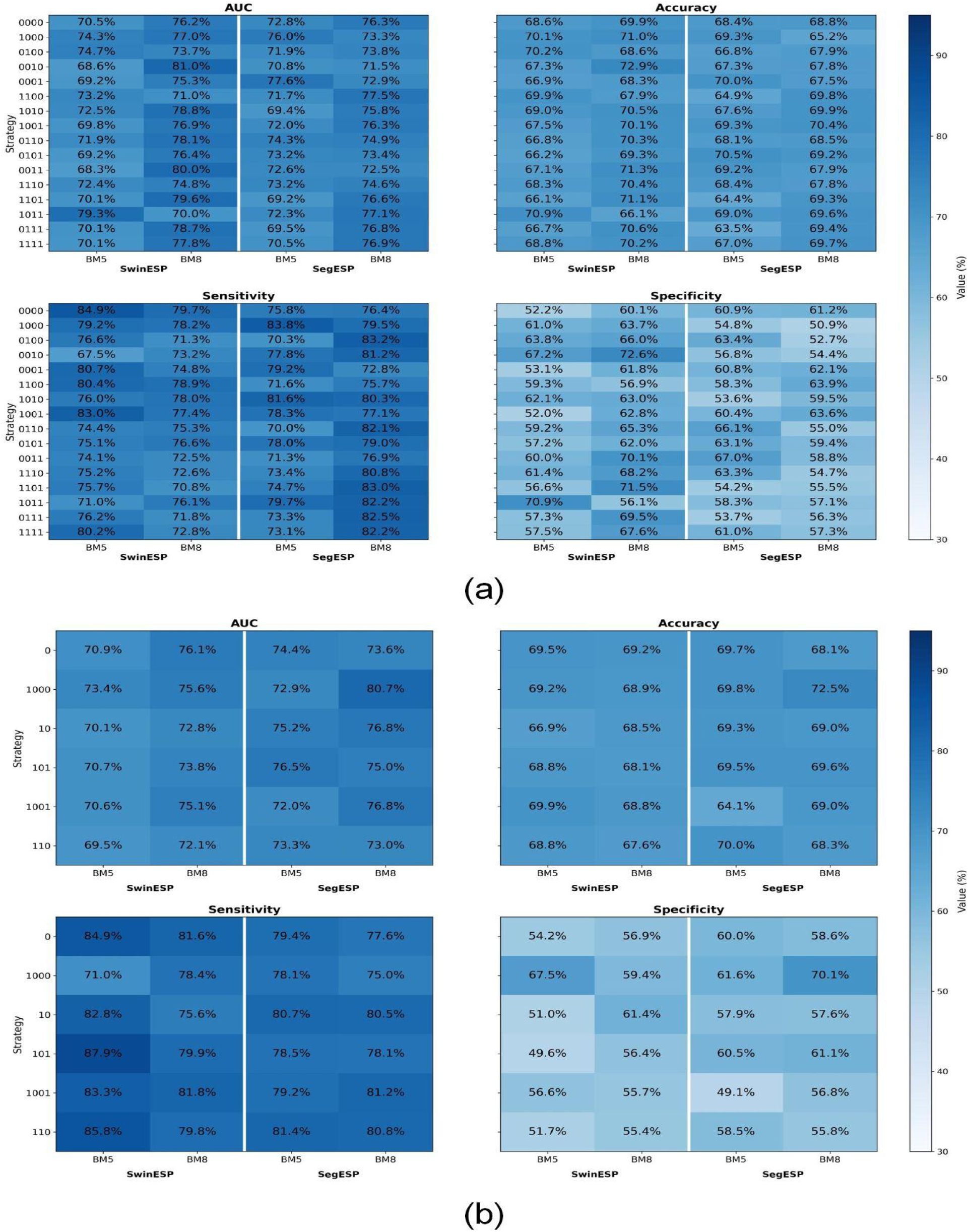
AUC, Accuracy, Sensitivity and Specificity heatmaps for stages 1 and 2. (a) Heatmap for all 16 strategies with BM5 and BM8 with SegESP and SwinESP. (b) Heatmap for the six selected strategies with BM5 and BM8 with SegESP and SwinESP.

**Figure 6.**
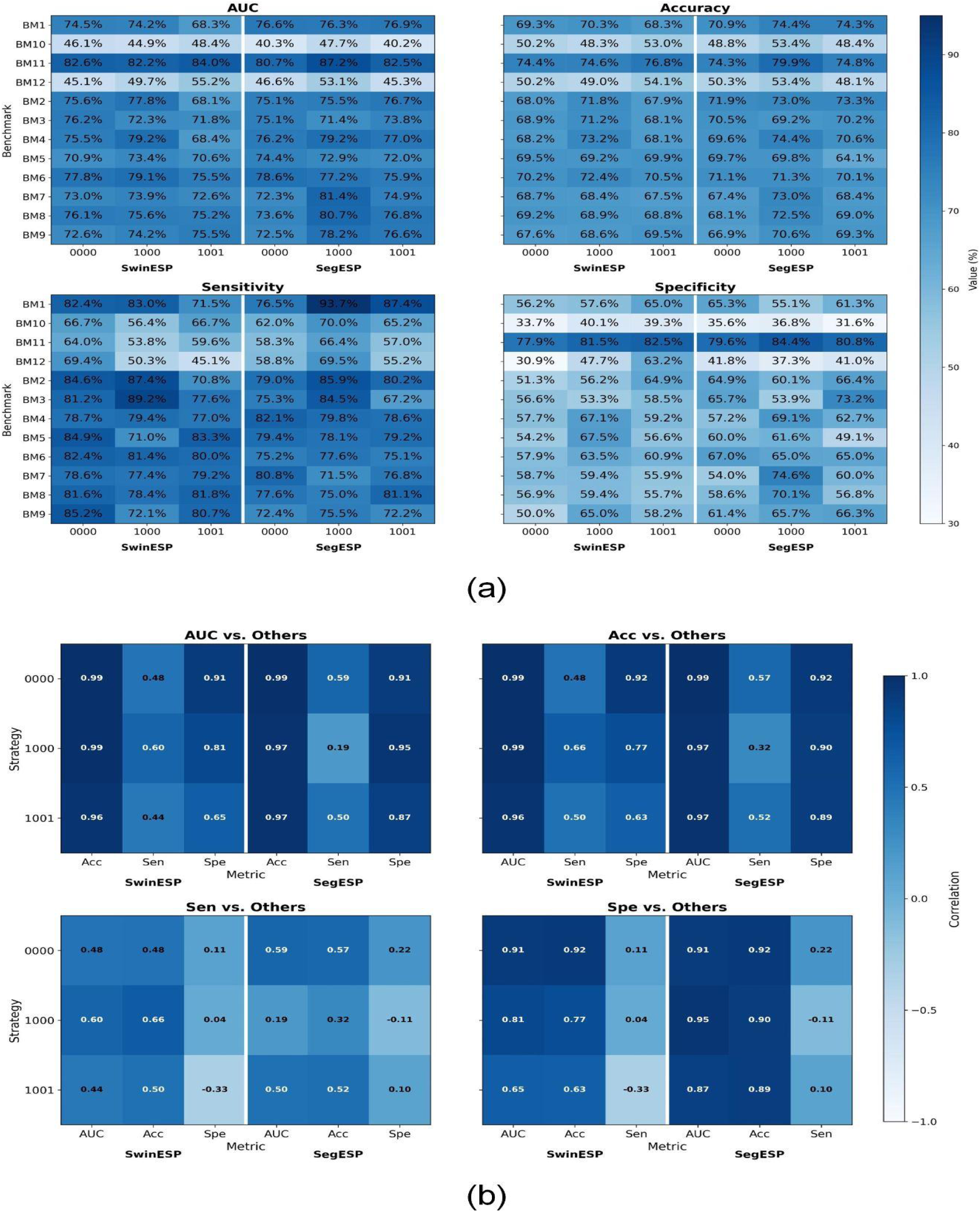
AUC, Accuracy, Sensitivity and Specificity heatmaps for stages 3 with correlations. (a) Heatmap for all 12 benchmarks with three strategies. (b) Correlation heatmaps for all metrics.

Overall, CSeg-ESP models provide the best AUC scores for 50% of the benchmarks with each method providing the best AUC for three BMs each, CSeg-ESP-1 for BM7 and BM11-12 and CSeg-ESP-2 for BM2 and BM9-10. ViTESPOpt and ESPFOrmer each provide the best AUC scores for a quarter of the benchmarks each. ESPFormer gives the highest respective AUC scores of 79.8%, 82.7% and 80.4% for BM1 and BM4-5. The highest AUC scores of 76.2% and 79.1% respectively for BM3 and BM6 is provided by SwinOpt whereas SegOpt provides the highest score for only one BM, BM8 with 83.8%. In terms of other metrics, there are several BMs where other CViT-ESP architectures outperform CSeg-ESP-1. For example, the highest accuracy and accuracy for BM4 is provided by CSeg-ESP-2 at 73.1% and 80.8%, respectively, compared to respective values of 70.5% and 78.0% by CSeg-ESP-1 as shown in Table 4. Further, for BM10 (one of the tougher BMs), CSwin-ESP-1 has the highest accuracy of 70.9% due to it providing the best sensitivity of 64.3% for that BM compared to an accuracy of 68.7% and a sensitivity of 56.5% by CSeg-ESP-1. However, CSeg-ESP-1 has the best performance for several benchmarks for various metrics such as the highest sensitivities of 85.8% and 87.1% for BM1-2, the highest sensitivity of 63.4% for BM12 where other models do not perform well, the highest accuracies of 74.7% and 74.9% for BM7-8 and the highest specificities of 93.1% and 76.4% for BM11-12. Hence, this combined with consistently high metrics results in CSeg-ESP-1 providing the best performance.

### Results from comprehensive search-space analysis with hyperparameter selection

Figure 5 illustrates that the heat maps represent the different performance metrics achieved with all 16 strategies for the representative benchmarks (BM5 and BM8) for both architectures SegESP and SwinESP. Figure 5a demonstrates the AUC, accuracy, specificity and sensitivity in each quadrant in a clockwise direction starting from the top left for all strategies for both SwinESP and SegESP with BM5 and BM8. The strategies are represented on the y-axis labels as four binary digits corresponding to each of the four transformer blocks (akin to a flag bit representation). If the flag is ON or in other words, a particular binary digit is ‘1’, it represents a stage that was re-trained. For example, strategy 0100 indicates that the second transformer stage was re-trained. Recall that the input stage and the classifier head are tuned for all approaches.

Overall, we notice that strategies 1000 (first transformer stage fine-tuned), 1001 (first and fourth transformer blocks unfrozen and 0000 provide the highest AUC score though strategies 0000, 0000 and 0000 provide competitive AUC scores across all benchmarks. In general, even though some strategies such as 1101 and 0111 provide high AUC scores respectively up-to 79.7% and 78.7% with SegESP on BM8, they require re-training 3 transformer stages as opposed to strategy 0110 which has an AUC score that is lower up-to 0.7% but requires re-training only 2 transformer blocks. Hence, it was preferred over strategies that require tuning 3 transformer blocks. Similarly, 1001 which requires re-training 2 transformer blocks and has an added advantage of keeping the middle layers frozen, which is useful from the implementation perspective, provides an AUC score in excess of 76% with SegESP on both BM5 and BM8, but has a low AUC for both models on BM5, with SegESP providing an AUC of 72.0% on SwinESP providing an AUC of 69.8%. In contrast, strategy 1011 with fine-tunes an additional transformer block has similar numbers but also provides a 79.3% AUC with SwinESP on BM5. However, the sensitivity of ‘1001’ is 83% with SwinESP on BM5 and in excess of 77% on all other combinations of model and BMs. Hence, we select 1001 as opposed to 1011 as one of the strategies for the next stages. Further, other strategies such as 0000 that require no training of any transformer blocks (the original SwinESP and SegESP), give an AUC score in excess of 70% for all four BM and model combinations and accuracies in excess of 68%, which also demonstrates consistency. The AUC achieved by other strategies on either one of or on both BM5 and BM8 is about 10% lower while requiring considerable extra resources.

Based on this discussion, we shortlist the strategies 0000 (no transformer-block fine-tuning), 1000 (first transformer block), 0010 (third transformer block), 0110 (second and third transformer blocks), 1001 (first and last transformer blocks) and 0101 (second and last transformer blocks). In a similar fashion to Figure 5a, Figure 5b shows the achieved AUC, accuracy, specificity and sensitivity in each quadrant for all six shortlisted strategies for SegESP and SwinESP for both BM5 and BM8. Note that the metrics are averaged across all 10 hyperparameter sets used for these six strategies which counterintuitively makes the heatmap appear lighter. Judging based on the AUC score purely, the strategy of re-training only the first transformer block (‘1000’) provides the highest values with all AUC scores in excess of about 73% with the highest score of 80.7% on BM8 with SegESP; which also has the highest accuracy of 72.5%. Hence, we select this strategy over 0010 (re-training the third stage) which does provide very high sensitivity scores of in excess of 80% for three of the BM-model combos, but it has lower specificities leading to a lower accuracy for both BM5 and BM8 with SwinESP and SegESP. Additionally, we also select the original SwinESP and SegESP with no transformer blocks re-trained (strategy 0000) as it provides consistently high AUC scores and accuracy at a low computational cost. We choose one additional strategy 1001 among the strategies that require fine-tuning two transformer stages. The strategy 0100 is rejected due its consistently low AUC and accuracy scores including the lowest AUC of 69.5%. While 0101 slightly exceeds 1001 in performance, from the implementation perspective, the ability to fine-tune the first and last few layers is preferable as it is more conducive to deployment at the edge for real-time use with paradigms such as federated learning and split learning. Hence, we shortlist three strategies: 0000, 1000 and 1001.

### Results from all benchmarks with selected strategies

Figure 6 provides a comprehensive overview of the results achieved by training and validating both SwinESP and SegESP on all benchmarks using the final three shortlisted strategies. The best set of hyperparameters is used from stage two with the highest performing hyperparameters on BM5 assigned to BM1-6 and the best hyperparameters for BM8 assigned to BM7-12. A visual inspection of Figure 6a compared to Figure 5a demonstrates a significant improvement. Similar to Figure 5a, Figure 6a shows the AUC, accuracy, specificity and sensitivity in each quadrant. However, the y-axis now displays all BMs whereas the three shortlisted strategies for each architecture form the x-axis. In general, there is at least one strategy and architecture combination that provides an AUC score in excess of 78% for BM6-9 and 74% for BM1-5. All strategies struggle for BM10-12 and while the AUC is in excess of 80% for BM11 with all strategies for both SwinESP and SegESP, the prediction sensitivity is 66.4% or less. Despite specificities in excess of 77.9% which leads to accuracies of 74.4% or greater, models trained on BM10-12 have low predictive power. With the exception of BM5, there is at least one model and strategy combination that provides an accuracy in excess of 70% for all other BMs.

To select the best strategy for each benchmark, for each model we chose the strategy that provided the highest AUC score. Results show that the strategy of re-training only the first transformer stage was the most popular with 50% of the BMs having the highest AUC across SwinESP and SegESP. The strategies of re-training either no transformer stages or the first and last transformer blocks provided the highest AUC for 25% of BM-model combinations each. With 75% of the combinations requiring the re-training/fine-tuning of at least one transformer block, our approach to find the best re-training combination is justified. Hence, the selection of the best strategy of fine-tuning with its associated set of hyperparameters results in the outcome of SwinOPT and SegOpt. Because we are heavily relying on the AUC scores to make the selections, we study one more set of results to determine the implications of our approach.

In Figure 6b, we also compare how each metric correlates with other metrics for each strategy for both SwinESP and SegESP across all benchmarks. Because sensitivity is critical to avoid missed seizures but specificity is important in avoiding false alarms, it is not enough to analyze the AUC and accuracy. We want to ensure that the high accuracies translate to both high sensitivities and specificities. As this a class-balanced problem with an equal number of preictal and interictal samples (equal labels 0 and 1), we do expect the correlation to be low between the sensitivity and specificity. However, as long as we avoid negative correlations, it would indicate that the models and/or strategy offers a good trade-off between sensitivity and specificity and does not favor one outcome significantly more than the other. Overall, we see that the AUC heavily correlates with the accuracy for all three shortlisted strategies with both SwinESP and SegESP. However, there is a low correlation between the AUC and the sensitivity with the correlation being less than 0.6 for SwinESP and less than 0.5 for SegESP. Ironically, this is because the sensitivity of SegOpt and SwinOpt is relatively high. Unfortunately, correlations of up-to 0.92 between the accuracy and specificity indicate that low specificity values (as low as 33.7%) bring down the accuracy. This is reinforced by several strategies exhibiting negative correlations between the sensitivity and specificity such as -0.33 for SwinESP with strategy 1001 and -0.11 for SegESP with strategy 1000. This shows that both SegOpt and SwinOpt may struggle to offer a good trade-off between sensitivity and specificity while maintaining a high accuracy. Hence, SegOpt has the highest sensitivity 75.8% but a relatively low accuracy of 69.8%.

### Complexity analysis

Figure 7 illustrates the complexity in terms of the number of training experiments needed by our proposed strategy under different conditions compared to an exhaustive search or ‘brute force’ approach. Figures 7a and 7b plot the complexity for our scenario where there are *L* = 2 architectures, each comprising *n* = 4 transformer blocks, that need to be trained for *D* = 12 datasets where *D*’ = 2 representative datasets are chosen. The models are tuned over a set of *H* = 10 hyperparameters. Figure 3a shows a linear plot restricted to between *n* = 1 and *n* = 4 whereas Figure 3b shows a log-scale plot up-to *n* = 20. Recall that our proposed strategy comprises three steps where the number of experiments in the second step depend upon the number of strategies (*P*) selected based either on an objective or subjective measure. The value of *P* can be set as a function *g*(*n*) to simplify the analysis and procedure. Therefore, we plot the complexity of our proposed approach for different possible values of *g*(*n*); though we selected *P* = *n* log *n*. Notice from Figure 3a that for *n* = 4, a brute force approach would require approximately 3,800 experiments whereas our proposed approach requires around 500 training experiments in total. Overall, Figure 3b shows that as the number of transformer stages *n* is increased, our proposed methods would converge to a much lower number of needed experiments compared to an exhaustive search as long as *g*(*n*) is of polynomial complexity. In fact, this convergence is much faster if *g*(*n*) is of quadratic complexity or lower. Figure 3c how this effect is even more enhanced where there is an extraordinarily large number of experiments. For example, for architectures with 10 transformer blocks, an exhaustive search requires more than 1 billion experiments whereas our proposed method requires less than 1 million experiments (three orders of magnitude lower) if *g*(*n*) is of log, linear and log-linear complexity. For polynomial complexities (quadratic or cubic), the number of experiments is less than 10 million which is still two orders of magnitude lower compared to brute force. Overall, this demonstrates a clear capability of our proposed heuristic algorithm to discover suitable re-training combinations without requiring an exhaustive number of experiments. Figure 3d illustrates the worst-case scenario, where *L*, *D*, and *H* = 1, i.e. only one architecture is tuned for one-dataset with one set of hyperparameters. Even in that case, the proposed strategy converges to the brute force approach when the architecture contains more than 10 transformer blocks for most choices of *g*(*n*).

**Figure 7.**
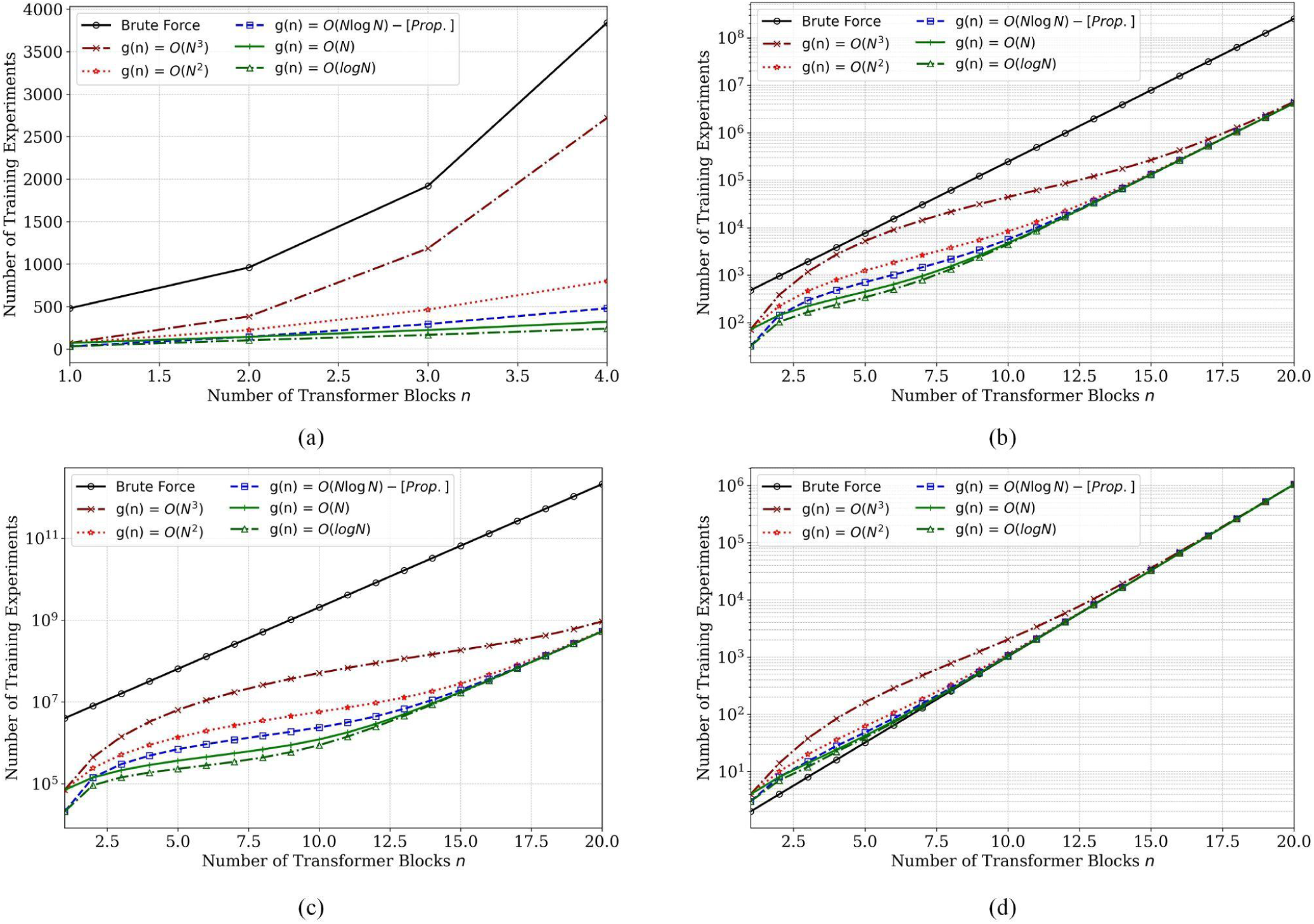
Complexity in terms of the number of Experiments. (a) Linear plot depicting the progression of the number of experiments as the transformer blocks are increased from 1 to 4 (our architectures have 4 blocks). (b) Log-scale plot depicting the progression under the same conditions as (a) but up-to 20 transformer blocks. (c) Improvement offered by our strategy in terms of reduced training experiments when there are a large number of architectures, datasets and hyperparameter sets that need to be tuned. (d) Impact of our strategy when there is no fine-tuning required and only the best combination of transformer blocks is sought.

### Comments on the performance of SwinOpt and SegOpt compared to ESPFormer

Observing the ROC curves for each benchmark in Fig. 4, we notice that while ESPFormer provides a better performance compared to both ESPSwinOpt and ESPSegOpt in BM1, BM2, BM4 and BM5, it provides a considerably lower performance from several other BMs including BM6 - BM9. Hence, the average AUC score across all BMs for ESPFormer is 69.0%, which was slightly better than SwinESP at 68.5% but slightly worse than SegESP at 69.7%. Applying our proposed strategy to find the best retraining combination of transformer blocks significantly enhances the performance in terms of the AUC score. For example, SwinESPOpt provides an AUC score of 70.8% which is an improvement of 1.8% compared to ESPFormer and 2.3% compared to SwinESP. Moreover, SegESPOpt even has a higher performance with an AUC score of 72.2% which is an improvement of 3.2% compared to ESPFormer and 2.5% compared to SegESP.

### Additional experimentation and analysis

#### Inference on CHB-MIT with CViT-ESP

Figure 8 summarizes the inference results on the CHB-MIT dataset, offering a comprehensive view of benchmark distribution, metric relationships, and subject-specific performance. The two pie charts illustrate the proportional representation of different benchmarks in the dataset, with certain benchmarks such as BM1 and BM2 accounting for the largest shares. The correlation matrix highlights strong positive associations between accuracy and AUC, while sensitivity and specificity are only weakly correlated, indicating potential trade-offs between detecting seizures and minimizing false positives. The box plots of AUC and accuracy for individual seizures across subjects reveal substantial variability, with some subject-seizure pairs achieving high scores while others demonstrate wider ranges and lower medians. Bar plots further distill these findings, showing the best AUC and accuracy per subject, as well as average performance at the patient level. These results collectively suggest that while the model achieves strong performance for some subjects and patients, considerable heterogeneity remains across the cohort, emphasizing the ongoing need for personalized or adaptive approaches to seizure detection.

**Figure 8.**
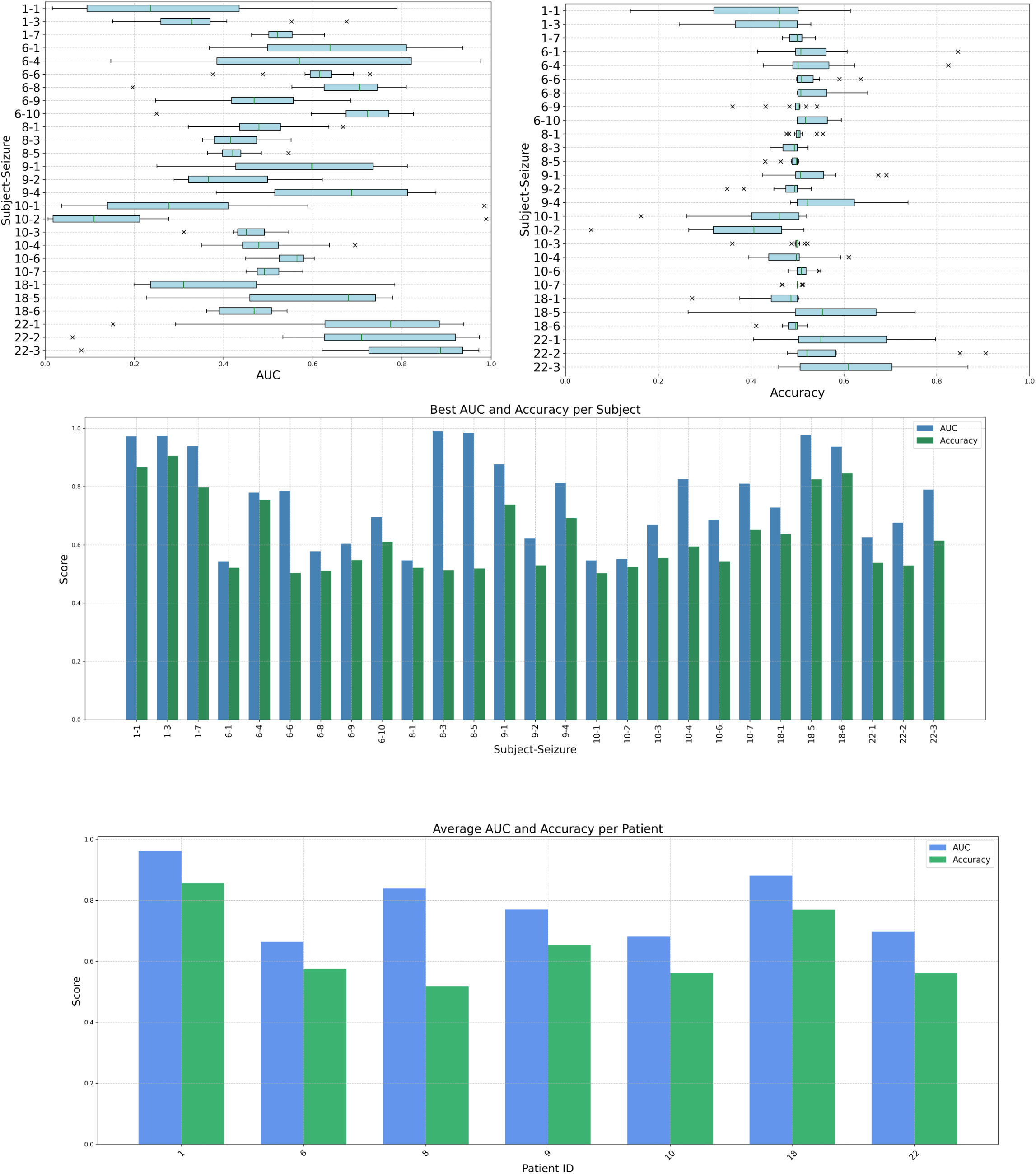
**Results of performing inference on the CHB-MIT dataset.**

## Discussion

Previous literature articulates the superiority of 1D CNNs and ResNets based on CNNs for time-series classification tasks. Although our CViT-ESP-1 and CViT-ESP-2 architectures only differ in implementation, the effects are clear. For example, as Tables 1 and 3 show, compared to CSeg-ESP-2, the average accuracy of CSeg-ESP-1 across all BMs is 1.7% higher, the sensitivity is 3.6%, a higher specificity by 2.3% and the AUC is also better by 1.3%. Similarly, CSwin-ESP-1 beats CSwin-ESP-1 in AUC, accuracy and sensitivity respectively by 2.2%, 3.1%, and 6.5%. The only outlier is that the specificity of both is 70.8%. In general, CViT-ESP architecture provides the best performance for the higher BMs including all of BM7-12. Particularly, the performance on BM10 and BM12 is comparable to other BMs with CViT-ESP where all other models failed. One potential reason is that although these benchmarks have a smaller number of subjects and seizures, they include more preictal data due to a longer SPH, which allows the combined CNN-ViTs to better extract preictal patterns and biomarkers. For example, BM10 has 72 seizures as opposed to BM2 which has around 823 seizures but uses a preictal time of 30 minutes compared to 5 minutes in BM4. While the number of seizures is reduced ten-folds, the number of preictal and interictal segments only reduce by 50% (approximately 100k in BM4 versus around 50k in BM10) while the number of samples available per seizure increases 6x. Further, we discover that architectures based on the SegFormer perform better compared to Swin-based architectures for the seizure prediction task. Table 1 shows that with respect to the average AUC score across all BMs, SegESP beats SwinESP by 1.2%, SegOpt beats SwinOpt by 1.8%, CSeg-ESP-1 beats CSwin-ESP-1 by 1.0% and CSeg-ESP-2 beats CSwin-ESP-2 by 1.9%. There may be several potential reasons for this including the addition of normalization layers in every transformer stage for better stability, a balanced number of transformer blocks in every transformer stage and the use of GELU activation which probably models a chaotic time-series such as EEG better. Further, while SwinESP architectures apply scaled cosine when calculating attention scores, they use traditional embeddings as opposed to the learnable embeddings in SegESP architectures. As our results with the lightweight ESPFormer also demonstrate, using learnable encodings and embeddings improves the performance of transformer-based architectures in classification tasks. Recall that SegOpt provided the best sensitivity of 75.6% but with a low specificity 63.4%.Whereas this represents superior predictive power, it will also lead to the generation of many false alarms. In contrast, though CSeg-ESP-1 has lower predictive power with a sensitivity of 70.8%, it will also have the propensity to generate less false positives with a 10.0% higher specificity leading to a 2.7% better accuracy. This also offers a trade-off where subjects that suffer from frequent seizures can use a more sensitive model while subjects with less frequent seizures may use a more specific model. Moreover, the accuracy of CSeg-ESP-1 is lower than the AUC by 4.9% which indicates that there is significant room for improvement with techniques such as setting variable thresholds. For example, in our work because the problem is class-balanced, we assign all segments with a score greater than 0.5 to the label ‘1’, this can be adjusted based on metrics like the weighted F-1 score for different benchmarks and subjects.

### Limitations

There are a few limitations in this work which conversely offer interesting future directions. CViT-ESP design currently does not consider multiple CNN configurations and was not rigorously cross-validated with different hyperparameters. Further, the proposed heuristic algorithm for fine-tuning SegOpt and SwinOpt can be combined with CViT-ESP, as in the current version, all transformer stages are frozen during training. Lastly, the CViT-ESP-2 version only differs superficially from CViT-ESP-1. While previous results support the use of 1D-CNNs, an interesting experiment would be to combine actual 2D-CNNs with pre-trained ViTs. An example configuration for a four-stage CNN with ma pooling could be: 1280 × 20 × 512 → 1280 × 20 × 256 → [640 × 10 × 256] where the first two terms are convolutions and third term represents max pooling, followed by: 640 × 10 × 128 → 640 × 10 × 64 → [320 × 5 × 64] with the a 320 × 320 representation being the 2D flattened version suitable for ViTs. For SegOpt and SwinOpt, though the proposed algorithm combines objective and subjective approaches, the purely objective version needs to be tested without subjective measures to demonstrate theoretical limits. Most importantly, the first step of the algorithm still has exponential complexity which needs a new innovative optimization method to be reduced at least to a polynomial complexity in terms of the experiments required. Lastly, instead of shortlisting a specific number of strategies in steps 2 and 3, they can be selected based on a cut-off validation metric. The caveat is that it will be hard to derive a theoretical bound on the complexity and will need a few training experiments for different architectures, tasks and datasets.

## Conclusions

In this paper, we designed, developed and then demonstrated that a clear progression in predictive performance is achievable through the strategic adaptation of pre-trained Vision Transformers (ViTs) for EEG-based seizure prediction. In this paper, we showcased the evolution of the model design including simple transfer learning with ResNets (for comparison) to sophisticated custom architectures that are integrated with pre-trained vision transformer models. Our findings highlight three major advancements: (1) the CVIT-ESP family of architectures, which integrates custom N-dimensional CNN stages with pre-trained ViTs, consistently outperformed existing methods, achieving a peak average Area Under the Curve (AUC) of 76.4%. In addition, we also presented the design and performance estimation of a lightweight, custom-designed transformer specifically engineered to mitigate overfitting on limited-scale EEG datasets, called ESPFormer. Furthermore, we developed a heuristic strategy to find the optimal fine-tuning combination of transformer blocks to be trained for achieving highest performance. The validation of all of the strategies was accomplished using patient-independent MLSPred-Bench, which is a combination of 12 different benchmarks with various seizure prediction horizons (SPH) and the seizure occurrence period (SOP). This ensured that our models are evaluated on a number of real-world scenarios, enhancing model generalizability and providing a more robust balance between sensitivity and specificity. Collectively, these results suggest that leveraging pre-trained vision models - when combined with domain-specific CNN front-ends and optimized re-training strategies - offers a resource-efficient path toward clinical-grade, patient-independent seizure prediction systems.

## Acknowledgments

Primary funding for this project is provided by NSF OAC-2312599 (FS). The content is solely the responsibility of the authors and does not necessarily represent the official views of the National Science Foundation.

## Competing Interests

Dr. Fahad Saeed is the Founder of AI-NeoTech LLC which is a startup that develops ML/AI solutions for diagnosing, characterizing and predicting various neuro disorders including epilepsy. All the remaining authors declare no conflict of interest.

## Author Contributions

FS, AAK and SC conceived and designed the project. FS and AAK acquired the MS and fMRI data. FS, AAK and SC analyzed the data, and designed the experiments. SC wrote the code and scripts and pipelines. FS, AAK and SC wrote the paper and all authors contributed to the revisions and approved the final version. FS contributed to all aspects of the project. AAK and SC contributed equally to this work.

## Notes

### Competing Interest Statement

Fahad Saeed is a Founder of startup AI-NeoTech LLC which is a company that develops AI technology for various health and biomedical data including epileptic seizures.

